# Clofarabine commandeers the RNR-α—ZRANB3 nuclear signaling axis

**DOI:** 10.1101/830745

**Authors:** Marcus J. C. Long, Yi Zhao, Yimon Aye

## Abstract

Ribonucleotide reductase (RNR) is an essential enzyme in DNA-biogenesis and a target of several chemotherapeutics. Here we investigate how anti-leukemic drugs [e.g., clofarabine (ClF)] that target one of the two subunits of RNR, RNR-α, affect non-canonical RNR-α functions. We discovered that these clinically-approved RNR-inhibiting dATP-analogs inhibit growth by also targeting ZRANB3—a newly-identified DNA-synthesis promoter and nuclear-localized interactor of RNR-α. Remarkably, in early time points following drug treatment, ZRANB3-targeting accounted for most of the drug-induced DNA-synthesis suppression and multiple cell types featuring ZRANB3-knockout/knockdown were resistant to these drugs. Additionally, ZRANB3 plays a major role in regulating tumor-invasion and H-*ras*^G12V^-promoted transformation in a manner dependent on the recently-discovered interactome of RNR-α involving select cytosolic-/nuclear-localized protein-players. The H-*ras*^G12V^-promoted transformation—which we show requires ZRANB3-supported DNA-synthesis—was efficiently suppressed by ClF. Such overlooked mechanisms-of-action of approved drugs and a new example of non-oncogene addiction, which is suppressed by RNR-α, may advance cancer interventions.

---

Ribonucleotide reductase (RNR) is a dual-subunit enzyme essential for dNTP synthesis and maintaining absolute and relative dNTP levels^1–3^. One subunit, RNR-β, is present only in S-phase. The other subunit, RNR-α, is present throughout the cell cycle^4^. RNR-reductase activity occurs by dint of a catalytically-essential cysteine (C429) within RNR-α that is activated via a proton-coupled electron transfer process initiated by a metallocofactor within RNR-β^5^. An alternative RNR-β-subunit, RNR-p53β, is constitutively present but exists only at low levels^4^. Because both proof-reading and mis-incorporation rates are acutely sensitive to relative and absolute dNTP levels^6^, dNTP-pool maintenance regulated by RNR— beyond its role in dNTP provision—is essential for genome integrity^7^. RNR-α is an excellent sensor of (d)NTPs. This (deoxy)nucleotide-sensing function is coupled to reductase activity: ATP is an allosteric promoter of RNR-enzyme activity; dATP is an allosteric downregulator^1^. The dATP-driven downregulation of RNR activity is functionally coupled to RNR-α-specific hexamerization^2, 8–10^. Several FDA-approved dA-mimetic nucleoside antimetabolites, e.g., clofarabine (ClF) (**Figure 1**), also cause RNR-α hexamerization^11, 12^. RNR-α oligomers (endogenous and ectopic) have been isolated from HEK293T, HeLa, and COS-7 cells post ClF exposure^13^. Electron microscopy showed hexameric species similar to those formed with recombinant RNR-α in vitro^13, 14^. Not unexpectedly, nucleoside metabolites such as ClF, which is FDA-approved to treat leukemias but is not clinically-used against solid tumors, elicit numerous bioactivities, including misincorporation, DNA-damage, apoptotic signaling, and possibly affecting unfolded protein response^15–19^. However, expression of RNR-α(D57N), a hexamerization-defective-but-reductase-active mutant resistant to ClF, abrogates ClF-induced cytotoxicity^12^. Conversely, gemcitabine (F2C) (a nucleoside antimetabolite approved to treat pancreatic cancer, which functions as a mechanism-based inactivator of RNR-α^5^ but does not cause RNR-α-specific hexamerization)^12, 20, 21^ and triapine (3-AP) (that targets RNR-β)^22^ (**Figure S1A**) are both only weakly protected by RNR-α(D57N) expression^12^. Thus, RNR-α is a key early target in the pharmaceutical profile of RNR-inhibiting nucleoside-antimetabolite drugs, and in the case of ClF, RNR-α-hexamer induction is required for bioactivity.

**Figure 1.**
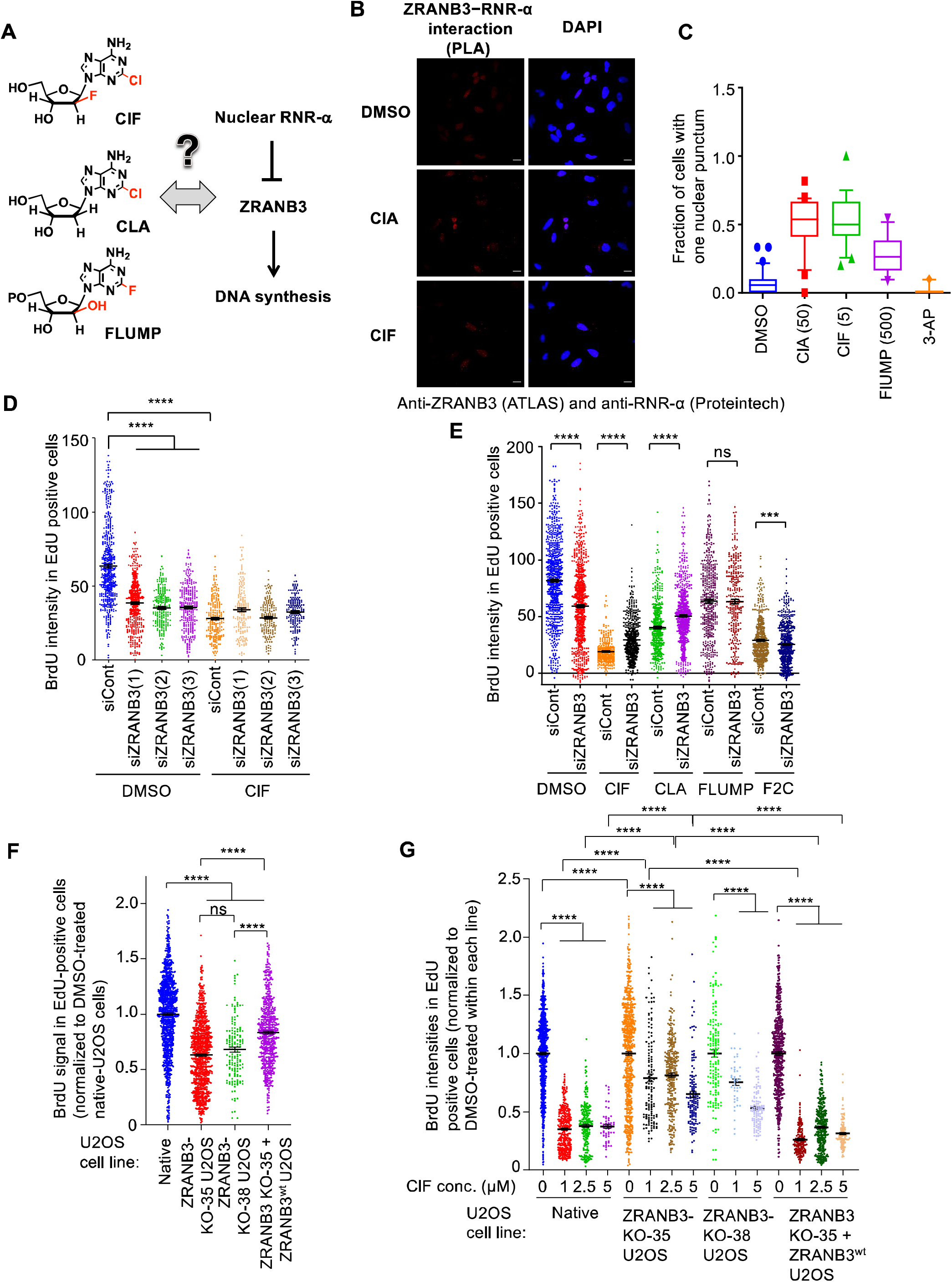
Early DNA-synthesis suppression driven by antileukemic nucleotides ClF/CLA/FLUMP is dependent on ZRANB3. (Relates to **Figure S1–S6**). **A.** This study for the first time interrogates the potential interplay between clofarabine (ClF), cladribine (CLA), and fludarabine monophosphate (FLUMP) and the recently-discovered ZRANB3—RNR-α signaling axis (also see **Figure S1B**). **B.** Representative images of PLA showing HeLa cells treated with DMSO, CLA (50 µM) and ClF (5 µM). Scale bar, 10 µm. **C.** Quantitation of cells with one or more punctum in nucleus (for two or more puncta per nucleus, see also **Figure S1E**). Concentrations (µM) used are indicated in parentheses. **D.** HEK293T cells were treated with the indicated siRNA for 2 days then they were either treated with DMSO or ClF (5 µM) for 30 min. After this time, dual-pulse labeling was carried out with a pulsing time of 30 min for each dT analog (EdU, to clearly label S-phase cells, followed immediately by BrdU, which was used to quantify DNA-synthesis rate during the second pulse). After this time, cells were washed with PBS, then fixed with methanol. Data were analyzed by confocal imaging and density of BrdU signal in EdU positive cells was measured. **E.** Similar to **D**, but cells were treated with the indicated compound (F2C and ClF 5 µM, CLA 50 µM, FLUMP 200 µM) for 30 min in each case. Note: Figures D and E are derived from separate batches of cells from different passage numbers treated on different days. Some variation is observed between runs. Also see **Figure S1F**, and **S2A–B**. **F.** U2OS, ZRANB3-KO-35 U2OS, ZRANB3-KO-38 U2OS, and ZRANB3-KO-35 U2OS cells expressing ZRANB3^wt^ were treated with DMSO for 30 min. Afterwards, dual-pulse labeling was carried out with a pulsing time of 30 min for each dT analog (EdU, to clearly label S-phase cells, followed immediately by BrdU, which was used to quantify DNA synthesis during the second pulse). Cells were then washed with PBS, fixed with methanol followed by DPA. Data were analyzed by confocal imaging and density of BrdU signal in EdU positive cells was measured. Also, see **Figure S4A–C and S5A–B**. **G.** Similar to **F**, but the indicated U2OS lines were treated with DMSO or different concentrations of ClF for 30 min. Note: the DMSO data in **G** are recapitulated from **F**.

RNR-α is linked with numerous unexpected activities, including tumor suppression, for which there is little link to RNR’s canonical reductase activity^2^. These data hint at non-canonical roles for this protein. Interestingly, knockdown of RNR-α *promotes* tumor growth and transformation^23^, indicating that non-canonical activities of RNR-α may be, in some instances, dominant over the need to supply nucleotides. However, the necessity of RNR-α for dNTP provision means that correlating RNR-α-levels and expression in tumors is overall a complex process. Nonetheless, there is some evidence that RNR-α-expression is positively correlated with remission and survival^2^, for which there remains little explanation. Surprisingly, little basic science work has set about understanding these phenomena.

Change in protein quaternary state is an established means by which many enzymes acquire moonlighting functions^24^. We thus hypothesized that upon hexamerization, RNR-α gains ancillary functions independent of its role in ribonucleotide reduction. We recently discovered that hexameric RNR-α translocates into the nucleus within ∼30 min following cell stimulation by hexamer-inducers, e.g., dA and ClF (**Figure S1B**). Nuclear RNR-α—regardless of oligomerization state—binds ZRANB3^25^, a poorly-characterized protein that was primarily associated with the DNA damage response (DDR)^26, 27^. In the canonical model of ZRANB3-biology, following DNA damage, ZRANB3 associates with ubiquitinated PCNA to coordinate template switching to reinitiate stalled replication forks and bypass DNA lesions^28–30^; however, we found no indication of DDR-upregulation in our studies^25^. We further found that in cells subjected to DNA damage, the newly-identified nuclear-RNR-α—ZRANB3 interaction is ablated. This result implies that nuclear RNR-α regulates an overlooked ZRANB3-function occurring exclusively in non-damaged cells. We subsequently showed that ZRANB3 promotes DNA synthesis and passage into S-phase. Our findings agree with initial reports that ZRANB3 and PCNA interact in the absence of DNA damage, although the function of this interaction was unknown^28^. Specifically, we found that knockdown of ZRANB3 elicits 30-40% decrease in DNA-synthesis rates in S-phase cells^25^. The basal function of ZRANB3 requires PCNA association, and nuclear RNR-α binds ZRANB3 competitively with PCNA. Indeed, the ZRANB3—nuclear-RNR-α interaction occurs through the APIM domain of ZRANB3, one of the two functionally-coupled PCNA-binding sites within ZRANB3.

However, many key questions underlying the newly-defined RNR-α-interactome remain unanswered, both in terms of the RNR-α—ZRANB3 signaling axis and the role of ZRANB3 in promoting DNA synthesis in non-damaged cells. For instance, the nuclear content of RNR-α is increased by ClF. Nuclear RNR-α inhibits the DNA-synthesis-promoting function of ZRANB3. But ZRANB3 is required for rapid proliferation, which is a hallmark of cancer cells. Thus, one major therapeutically-relevant question constitutes the extent to which ZRANB3 plays a role in the clinical action of ClF (**Figure 1A**). Our previous studies^25^ explored this link using dA. However, dA is highly pleiotropic and acutely toxic, and must be administered at high (mM) concentrations. Such concerns raise questions concerning the role of adverse/secondary effects following dA stimulation that are not easily parsed from on-target effects. Here, we exploit the principle of genetic epistasis^31^, classic knockdown/knockout/rescue, and drug treatment to directly establish the involvement of ZRANB3 in ClF bioactivity in several independent lines. These questions are particularly interesting as ClF, at low (nM–µM) concentrations, induces rapid and persistent RNR-α-hexamerization^11, 13^ and consequent nuclear translocation^25^ in various cell types. As target occupancy by drugs in vivo is typically moderate especially when clearance is taken into account^32^, this mechanism could be important in the pharmaceutical programs choreographed by ClF and other RNR-α-hexamer-inducing anti-leukemic/-lymphoma drugs^12^, e.g., cladribine (CLA) and fludarabine-monophosphate (FLUMP) (**Figure 1A**). Such evidence would add weight to the growing data showing the importance of RNR-α translocation in ZRANB3-regulation (**Figure S1B**). By extension, such evidence would also further highlight the importance of the unappreciated role of ZRANB3 modulating *basal* DNA-synthesis rates (i.e., in cells not treated with DNA-damaging agents).

We were equally curious about how the unexplained tumor suppression roles of RNR-α^2^ could be related to the ZRANB3-dependent signaling axis for DNA-synthesis promotion in non-stressed cells (**Figure S1B**). In this regard, it is paramount to understand the roles of nuclear RNR-α and ZRANB3 in tumor suppression and how these roles are linked. Here we investigate how these nucleoside antimetabolites intersect with the ZRANB3/nuclear-RNR-α signaling axis (**Figure 1A**), using ZRANB3 knockdown- and knockout cells and nuclear-RNR-α-overexpression. Our data document that ZRANB3 function is indirectly modulated by ClF and its variants in HEK293T, U2OS, and NIH-3T3 cells, via RNR-α-hexamerization-dependent RNR-α-nuclear signaling. Our further experiments indicate that ZRANB3 is required for H-*ras*^G12V^-induced focus formation, which we demonstrate is suppressed by ClF.

## Proximity ligation assay confirms the association of endogenous ZRANB3 and endogenous RNR-α in cells

We have shown the ZRANB3—RNR-α interaction: directly, through immunoprecipitation (IP); indirectly, through the loss of ectopic ZRANB3—PCNA interaction by both IP and puncta-formation using immunofluorescence (IF)^25^. We thus first validated endogenous ZRANB3—RNR-α association in an in-cell assay. We used the proximity ligation assay (PLA)^33^. Although PLA is not a direct measurement of association, it is locale-specific and it can analyze endogenous proteins, with no need for epitope-tagging or overexpression in cells. PLA requires two antibodies of different species that (1) detect endogenous proteins and (2) are IF compatible. We previously disclosed RNR-α-antibodies from both mouse and rabbit that function well in IF and western blot (WB)^13, 25^. However, we have had difficulty validating a ZRANB3-antibody that functions *in IF*^25^: our previous antibody that showed, among several non-specific bands, one specific ZRANB3 band by WB (Bethyl laboratories), gave almost-exclusively-cytosolic staining by IF (**Figure S1C**). We here identified a rabbit-anti ZRANB3 antibody (ATLAS HPA035234) that functioned sufficiently well for IF: nuclear signal observed from this antibody was reduced around 70% by three different ZRANB3-specific siRNAs, relative to a non-targeted siControl (**Figure S1D**) (note: this result further validates the ZRANB3-siRNAs used in this work). WB analysis showed similar knockdown-efficiency. We conclude that the nuclear signal from this ATLAS antibody is due to a specific interaction with ZRANB3. Notably, this antibody also shows cytosolic staining that is *not* decreased following siZRANB3-treatment (**Figure S1D**). This non-specific cytosolic staining does not bias our analysis of the ZRANB3 nuclear-specific protein.

We proceeded to the PLA using the ATLAS anti-ZRANB3-/mouse anti-RNR-α-antibody combination. DMSO treatment gave weak background staining of the nucleus with some elevated signal in the nucleolar region in HeLa cells (**Figure 1B**). A few cells showed nuclear puncta, likely reflecting that ZRANB3 and RNR-α interact without stimulation. Treatment of cells with nuclear-translocation inducers (ClF, CLA, FLUMP^25^) increased the number of nuclear puncta observed and increased the number of cells with more than one nuclear punctum (**Figure S1E**). 3-AP—an RNR inhibitor that does not elicit RNR-α-hexamerization/-translocation^22^ (**Figure S1A**)—did not increase nuclear puncta (**Figure 1C, and S1E**). Unsurprisingly, as the anti-ZRANB3 antibody shows some cytosolic staining (**Figure S1D**) and the hexamer-inducers ClF/CLA/FLUMP alter the cytosolic interactome of RNR-α (namely, IRBIT and importin-a1) (**Figure S1B**) as well as increasing the nuclear proportion of RNR-α, we also identified (likely-RNR-α-dependent/ZRANB3-independent) increase in puncta in the cytosol. Since nuclear-ZRANB3 is the predominant source of signal derived from the ZRANB3-antibody (**Figure S1D**), the elevation of nuclear puncta that selectively occurs upon treatment only with multiple RNR-α-nuclear translocation-inducers (and not with 3-AP) is strongly consistent with our previous data.

## ZRANB3 is one of the earliest inhibition events following ClF treatment

We next investigated how ClF affects DNA synthesis in siRNA-knockdown control vs. siZRANB3-knockdown cells, using three different ZRANB3-targeting siRNAs [validated above by IF (**Figure S1D**), and previously, by WB^25^] in dual-pulse whole-nuclei staining assay (DPA). DPA was chosen because it gives a very sensitive measure of DNA synthesis, over a specific time period (allowing discounting of cells entering DNA synthesis during the assay, and giving more confidence that each signal measured is specific). In cells not treated with ClF, all three ZRANB3-siRNAs suppressed DNA-synthesis rates 40% relative to control siRNA (**Figure 1D**), consistent with our previous report^25^. ClF treatment alone led to an almost 50% suppression in DNA synthesis. However, when siZRANB3 cells were treated with ClF, DNA synthesis was not affected relative to siZRANB3 cells treated with DMSO in place of ClF (**Figure 1D**). Such reduced (or hypomorphic) response to ClF in siZRANB3 cells indicates that ZRANB3 is also inhibited (directly or indirectly) by ClF and that this inhibition *significantly contributes to the response seen in control cells*. This finding is similar to our previous report on cells stimulated with deoxyadenosine (dA)^25^. As DPA specifically measures DNA-synthesis rates in S-phase, it is not affected by changes in cell-cycle distribution or growth rates that we have reported in these ZRANB3-knockdown lines^25^, although changes in cell-cycle/growth are relatively minor compared to the observed, almost-completely-penetrative loss of response to ClF-treatment in ZRANB3-deficient cells. Finally, these data are also compelling as we employed siRNA- and drug-treatments, which give little scope for genetic compensation/selection that may occur during prolonged knockdown or knockout.

## ZRANB3 inhibition is an early event common to hexamer-inducing RNR-**α**-allosteric modulators

To further investigate the role of RNR-α translocation in this ZRANB3-mediated DNA-synthesis suppression process, we replicated these assays using different RNR-inhibiting nucleosides. Beyond ClF, we examined additional RNR-α-targeting reversible allosteric modulators^12^ that cause RNR-α-specific nuclear translocation^25^ (namely, CLA and FLUMP) (**Figure 1A**). Results from these compounds were compared to those from the irreversible mechanism-based inactivator, gemcitabine (F2C) (**Figure S1A**, a compound that does not induce either RNR-α-specific hexamerization or nuclear translocation, and that is also a frontline chemotherapeutic^2^). siZRANB3-treated cells showed resistance to all nucleoside drugs relative to siControl cells (**Figure 1E**). However, resistance was most apparent when the drug-treatment promoted RNR-α translocation, regardless of the magnitude of DNA-synthesis suppression.

Similar effects were observed upon treatment with natural hexamer-inducer dA (**Figure S2A–B**), with and without the co-treatment with non-hexamer-inducing dG/dC (**Figure S2B**). These results demonstrate that the observed rescue from ClF/CLA/FLUMP-induced replication-inhibition in siZRANB3-treated cells (**Figure 1D–E**) is an endogenous function, and not an artifact of drug treatment. We cannot rule out that some other factors contribute *weakly* to the observed hypomorphic responses to reductase-activity-downregulating drugs. However, these data indicate that RNR-α allosteric modulators—which elicit RNR-α-specific nuclear translocation (**Figure S1B**)—selectively target ZRANB3 as part of their pharmaceutical spectrum. The mechanism-based irreversible inactivator F2C (**Figure S1A**)—that does not elicit RNR-α-specific hexamerization^12^/translocation^25^—does not target ZRANB3. A similar effect was observed in NIH-3T3 cells expressing integrated shRNA-control or shZRANB3 (**Figure S1F**), thereby confirming that the effect is not cell-line-specific and is not unique to transformed cells.

To investigate this phenomenon further, we investigated the effect of ZRANB3-knockout (KO) and rescue in U2OS cells, a human osteosarcoma line that has been used extensively for DNA-damage research and specifically in studies involving ZRANB3 nuclear protein^28, 34^. We first validated that RNR-α-translocation occurred upon ClF-treatment in U2OS cells (3 µM, 30 min) (**Figure S2C**). We also validated the ZRANB3-antibody (ATLAS HPA035234) for use in IF in this line, by comparing ZRANB3-signal in U2OS cells, ZRANB3-KO-35 U2OS (nuclear signal depleted), and ZRANB3-KO-35 U2OS re-expressing ZRANB3^wt^ (nuclear signal restored, with a small percentage of cells hyper expressing ZRANB3, as is common in such lines) (**Figure S2D-E**). Similar outcomes in terms of ZRANB3-levels were also observed by WB (**Figure S2F**). With these key validations in hand, PLA assay was conducted in U2OS and a ClF-dependent increase in nuclear puncta was observed (**Figure S3A-C**). By contrast, in ZRANB3-KO-35 U2OS, a lower basal number of nuclear puncta was observed relative to U2OS; critically, there was *no* increase in nuclear puncta following ClF-treatment. Both basal and ClF-dependent nuclear-puncta formation was re-established in ZRANB3-KO-35 USO2 re-expressing ZRANB3.

Subsequent execution of DPA in two different ZRANB3-KO U2OS lines as expected showed suppressed DNA-synthesis in S-phase in non-stressed cells [**Figure 1F, S4A–C** (see DMSO-treated samples), and **S5A– B**]. Re-expression of wt-ZRANB3 in one of these KO lines rescued this basal DNA-synthesis rate [**Figure 1F** and **S4B** (images in top-row, left and middle panels)]. As predicted from PLA above, ZRANB3-KO-35 and -38 lines were also resistant to ClF-treatment for 30 minutes relative to U2OS (**Figure 1G**, and cf. **S4A** vs. **S4B**), and ZRANB3-KO U2OS-35 re-expressing ZRANB3^wt^ (**Figure 1G** and **S4B**: left vs. middle panels). ZRANB3- or RNR-α-expression was not affected by any of the different drug treatments in this line (**Figure S5A–C**), consistent with our previous data in other cell lines^12^. These data further show the generality of ZRANB3’s sensitivity to ClF, and underscore that although ZRANB3 is not absolutely required for DNA synthesis (since knockout cells can still synthesize DNA), this protein is an important adjuvant of DNA synthesis. Finally, although as noted above, knockout lines can show complications in terms of compensation effects, the ZRANB3-KO-35 U2OS cells re-expressing ZRANB3^wt^ were able to recapitulate effects observed in U2OS, indicating that a ZRANB3-specific effect is sufficient to explain these data.

We progressed to investigate how ZRANB3-knockdown affected late-stage susceptibility to the different drugs. For these assays, we used NIH-3T3 and HEK293T cells. The extent of cell proliferation (assessed by PI_50_ values, **Table 1**) for ClF, CLA, F2C, or 3-AP differed by no more than 1.7-fold between the control and knockdown cells, in both NIH-3T3 and HEK293T cells (**Figure S6**). Thus, ZRANB3-specific effects occur in the early time point and are not manifested at later stages following drug treatment, where a series of complex cytotoxic events involving DNA damage is known to ensue^15^. Critically, the ZRANB3—RNR-α interaction does not occur upon DNA damage^25^. Thus, the late-stage toxicity could readily override the effects of RNR-α-specific nuclear translocation that occurs rapidly and reaches saturation within 30 min at 5 µM ClF^25^. Prolonged treatment of cells with the drugs led to a non-detectable level of DNA synthesis, and thus, we are measuring a true reduction in DNA-synthesis rate, as opposed to a persistent DNA synthesis in the knockdown line (**Figure S7A**).

**Table 1.**
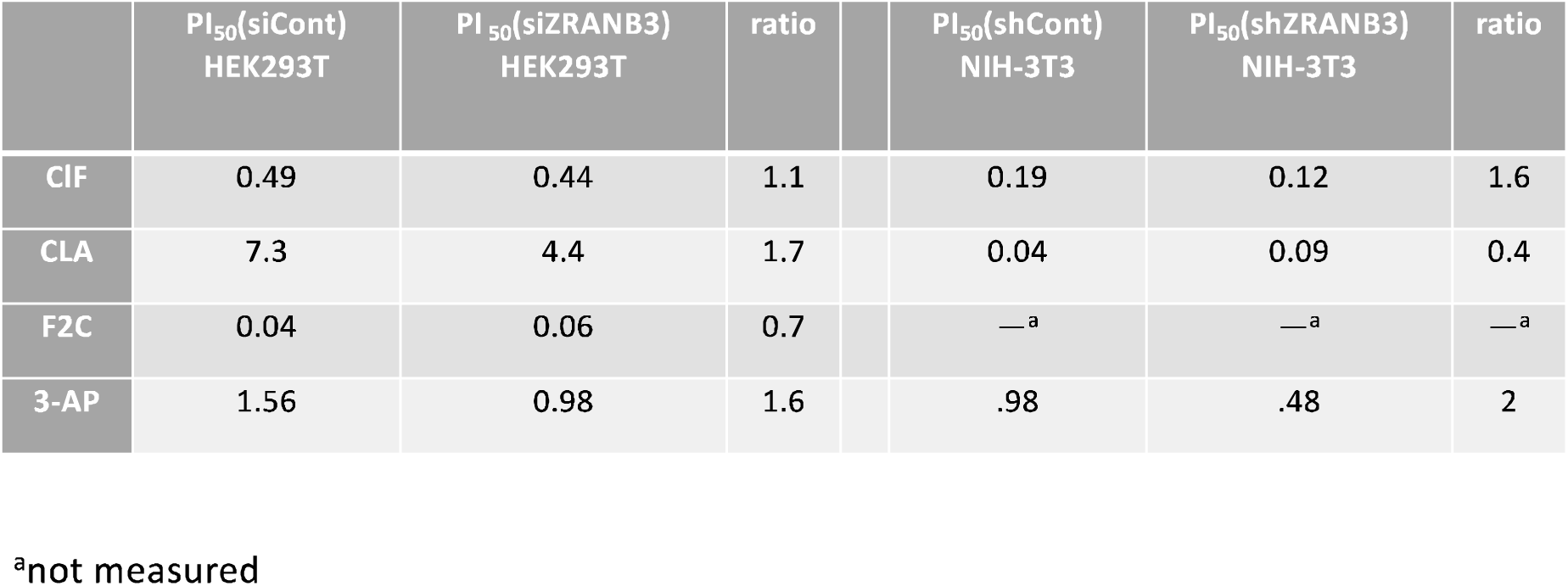
PI_50_ values for the indicated RNR-α inhibitors against ZRANB3-knockdown HEK293T and NIH-3T3 cells.

|  | PI<sub>50</sub>(siCont)<br>HEK293T | PI<sub>50</sub>(siZRANB3)<br>HEK293T | ratio |  | PI<sub>50</sub>(shCont)<br>NIH-3T3 | PI<sub>50</sub>(shZRANB3)<br>NIH-3T3 | ratio |
| --- | --- | --- | --- | --- | --- | --- | --- |
| CIF | 0.49 | 0.44 | 1.1 |  | 0.19 | 0.12 | 1.6 |
| CLA | 7.3 | 4.4 | 1.7 |  | 0.04 | 0.09 | 0.4 |
| F2C | 0.04 | 0.06 | 0.7 |  | —<sup>a</sup> | —<sup>a</sup> | —<sup>a</sup> |
| 3-AP | 1.56 | 0.98 | 1.6 |  | .98 | .48 | 2 |

<sup>a</sup>not measured

## RNR-α is required for clofarabine-induced ZRANB3 inhibition

We next examined responses to DNA synthesis upon combinations of siZRANB3, siRNR-α, and ClF treatment in HEK293T cells. siZRANB3 suppressed DNA synthesis. siRNR-α—that reduced both nuclear and cytosolic RNR-α levels (validated through IF, **Figure S7B–C**)—also suppressed DNA synthesis. However, siRNR-α lines did not show changes in DNA synthesis upon ZRANB3 knockdown (**Figure 2A and S7D–E**). This result indicates that RNR-α plays a role in the DNA-synthesis suppression that accompanies ZRANB3 knockdown. One simple explanation for such an observation is that loss of RNR-α increases basal ZRANB3 activity (i.e., DNA-synthesis promotion), rendering cells more able to sustain a partial loss of ZRANB3 following ZRANB3-knockdown (**Figure S7**).

**Figure 2.**
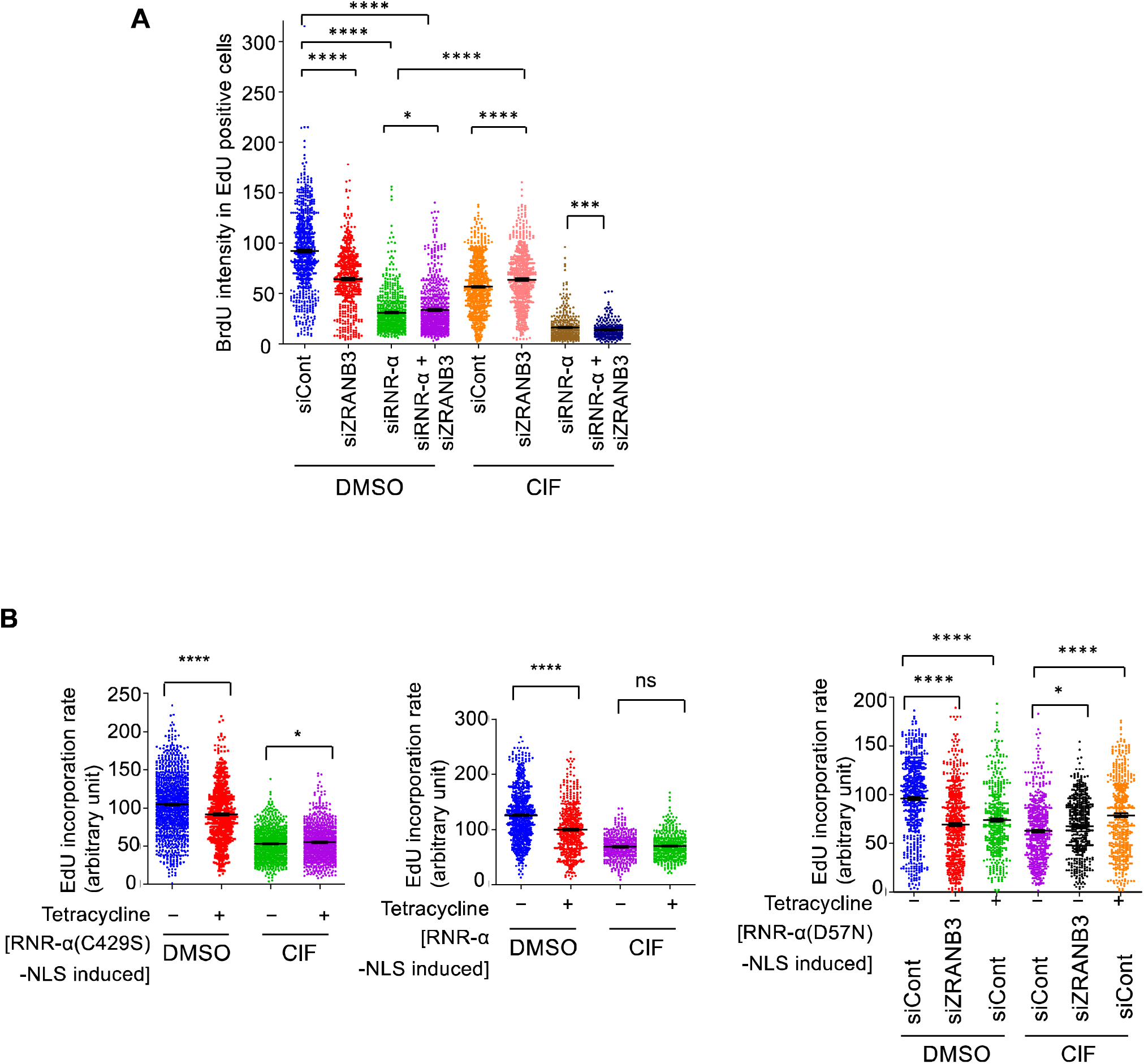
The response to ClF is dependent on RNR-α—independent of its reductase activity. (Relates to **Figure S7–S8**). **A.** HEK293T cells were treated with the indicated siRNA(s) for 2 days (note: for single siRNA treatments, an equal amount of siCont was added; for siCont alone treatment, 2X siCont was used), and subsequently, after 30-min treatment with ClF (5 µM) or DMSO, cells were subjected to dual-pulse labeling after which BrdU incorporation in EdU-positive cells was measured. **B.** HEK293T T-REx™ cells with single integrated copies of RNR-α (C429S)-NLS (*left*), RNR-α-NLS (*middle*), or RNR-α(D57N)-NLS (*right*) were exposed to the specific conditions (tetracycline exposure was for 2 d; ClF concentration was 5 µM; siRNA exposure was 2 days); EdU signal in EdU positive cells was recorded (performed this way because these cells have lower basal DNA synthesis, and are more susceptible to ClF than parent HEK293T cells).

We also investigated how ZRANB3-knockdown and RNR-α-knockdown affected response to ClF (**Figure 2A**). All lines, except the ZRANB3-knockdown line, showed downregulation in DNA synthesis following ClF treatment. The siRNR-α line showed moderate sensitization to ClF (and dA, **Figure S2A–B**) treatment. This observation may be expected given that these nucleosides also downregulate RNR-reductase activity (which is intrinsically linked to DNA-synthesis rate). However, DNA synthesis in the siZRANB3- and siRNR-α-treated cells was significantly reduced, relative to the ClF-treated siRNR-α-only lines. This rescue of sensitization to siZRANB3 in the presence of ClF could readily be explained as follows: knocking down RNR-α in the ZRANB3-knockdown line allows basal DNA synthesis to become ZRANB3-dependent [by reducing the extent of ZRANB3-inhibition by nuclear-RNR-α, since the amount of RNR-α in the nucleus is reduced (**Figure S7C, and S8**)]. This observation is also consistent with the fact that ZRANB3-knockdown does not promote DNA-synthesis suppression in the RNR-α-knockdown background (**Figure 2A**, DMSO-set). Thus, upon ClF treatment, translocated RNR-α meets a relatively low concentration of ZRANB3, which is contributing to DNA synthesis. The translocated RNR-α is sufficient to affect DNA synthesis in this double-knockdown background (**Figure 2A**, ClF-treated set, and **S8**).

## RNR-α nuclear accumulation suppresses ClF-induced DNA-synthesis downregulation regardless of RNR-reductase activity

To examine whether nuclear-specific RNR-α plays a role in ClF-induced DNA-synthesis inhibition, we used HEK293T T-REx™ monoclonal lines that upon tetracycline treatment express RNR-α bearing a nuclear localization tag (NLS). We have shown RNR-α-NLS (irrespective of wt/mutant) efficiently localizes to the nucleus, and inhibits ZRANB3’s DNA-synthesis promoting activity^25^. Expression of RNR-α-NLS regardless of reductase activity [wt, catalytically-active but hexamerization-and-nuclear-translocation-defective and ClF/dA-inhibition resistant mutant, RNR-α(D57N), or reductase-dead-but hexamerization-capable mutant, RNR-α(C429S)] diminishes DNA synthesis (**Figure 2B**). However, these RNR-α-NLS-expressing cells were resistant to ClF-induced DNA-synthesis inhibition, similarly to siZRANB3 lines (**Figure 2B** and **S2B**). Critically, this ClF-treatment-dependent suppression of DNA synthesis occurred regardless of RNR-reductase activity, and was not significantly more in the RNR-α(D57N) mutant that fully protects against ClF toxicity^12^ when the protein is cytosolic (**Figure 2B**). Thus, nuclear RNR-α suppresses the response to ClF regardless of RNR-reductase activity, ClF sensitivity, or RNR-α-hexamerization capability.

Furthermore, we have also shown that: (*1*) overexpression of RNR-α only weakly protects against late-stage toxicity of ClF, even though the drug-resistant point mutant [i.e., the reductase-active but hexamerization/translocation-defective RNR-α(D57N)] is highly resistant^12^; and (*2*) nuclear RNR-α-overexpressing cells are unable to inhibit DNA-synthesis in ZRANB3-deficient cells^25^, indicating that the nuclear role of RNR-α involves eliciting ZRANB3 loss-of-function. Taken together, we interpret these data to mean that ZRANB3 inhibition is an early event accompanying ClF treatment.

## ZRANB3-knockdown reduces wound healing in NIH-3T3 cells

We proposed that ZRANB3 inhibition could be a mechanism through which RNR-α overexpression elicits oncogene-induced tumor suppression, an unexplained observation made over the past two decades. ClF has a panoply of ascribed bioactivities, although at least in HEK293T T-REx™ cells, RNR-α inhibition is required for the majority of these responses. We thus asked the question: what phenotypic links are there between ClF/CLA/FLUMP treatment and RNR-α overexpression (a process that we have shown elevates nuclear RNR-α levels^25^)?

Answering this question led us to find two similar functional intersections between pharmacological action of RNR-inhibiting nucleos(t)ide drugs and genetic consequences of overexpressing RNR-α, namely, lower invasiveness and tumor suppression. It is worth noting that shRNA-Knockdown of ZRANB3 in most cell types, including NIH-3T3, proved challenging. In NIH-3T3, shZRANB3 was predominantly toxic. As alluded to above, we were able to establish one shZRANB3 line with reduced ZRANB3-protein levels. Beyond shControl line, we made use of shIRBIT cells. We previously showed that IRBIT-knockdown elevates nuclear RNR-α and displays phenotypes consistent with elevated nuclear RNR-α^25^ (**Figure S1B**). In wound-healing assays, we observed that both shZRANB3 and shIRBIT lines exhibited reduced wound closure compared to shControl lines (**Figure 3A**). This result is consistent with previous assays on RNR-α-overexpressing lines that showed reduced invasion in a Boyden chamber assay^23, 35–37^. These data demonstrate that ZRANB3-knockdown and IRBIT-knockdown show wound-closure/invasion-suppression phenotypes similar to the known behavior of RNR-α overexpression.

**Figure 3.**
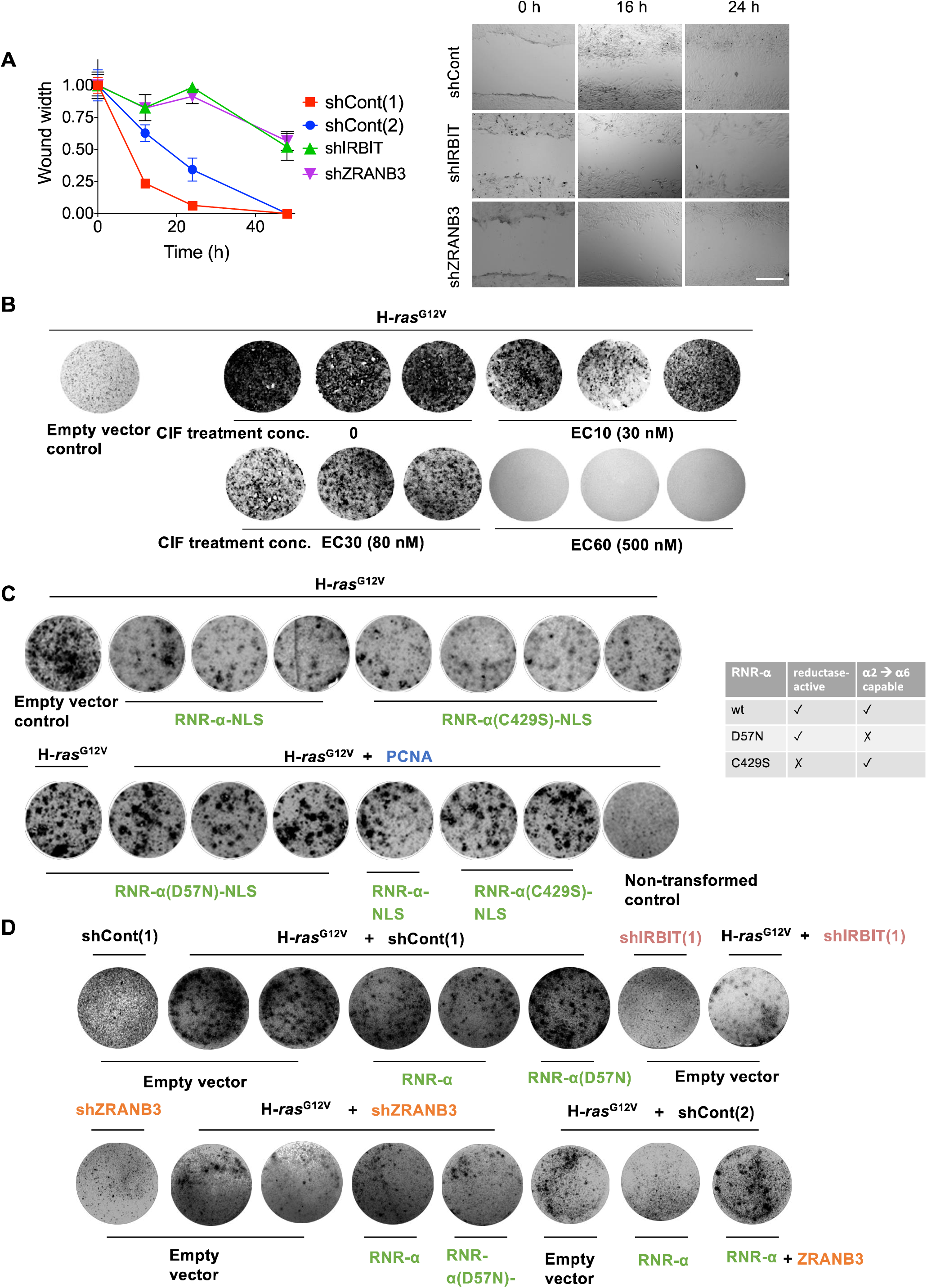
Phenotypes of ZRANB3-knockdown mirror those of RNR-α overexpression (which increases nuclear RNR-α levels). (Relates to **Figure S9**). **A.** NIH-3T3 cells expressing the indicated shRNA were grown on separate glass-backed plates each until a lawn of cells covered the whole plate surface. At this time, an autoclaved glass pipette was used to draw two scratches perpendicular to each other on each plate. The width of each scratch was measured as a function of time. Widths of each starting scratch were similar; quantitation (*left*) and representative images (*right*). Scale bar, 300 µm. **B.** NIH-3T3 cells were transfected with either H-*ras*^G12V^ or an empty vector plasmid before being treated with different concentrations of ClF (0, PI_10_ (30 nM), PI_30_ (80 nM), and PI_60_ (500 nM) as in **Figure S6** for shControl-1). The cells were cultured with ClF-supplemented medium, maintaining the same ClF concentration, for another two weeks before fixing with 2% paraformaldehyde in PBS, and foci were stained by crystal violet followed by water destaining. **C.** NIH-3T3 cells were either transfected with empty plasmid and transfection agent, or a plasmid expressing H-*ras*^G12V^, and the indicated RNR-α construct (Table on right shows wt vs. respective functional mutants employed), either empty vector and/or PCNA-GFP. Absence of GFP-PCNA was balanced using an empty vector. After 3 weeks, cells were analyzed as in **B**. **D.** Similar experiment to **C**, except NIH-3T3 cells expressing the stated shRNA were co-transfected with H-*ras*^G12V^. After 3 weeks, cells were analyzed as in **B**.

## ZRANB3-knockdown reduces focus formation in NIH-3T3 cells

We next investigated the role of ZRANB3/nuclear RNR-α in *ras*-induced transformation. Indeed, ClF/CLA/FLUMP have been previously shown to suppress anchorage-independent growth^38^. We first investigated how ClF affected focus formation in NIH-3T3 lines. Significant suppression of focus formation was observed even at the PI_10_ (concentration at 10% proliferation-inhibition) of the non-transformed lines, indicating that ClF is likely more effective against transformed cells than against non-transformed cells, similar to how ZRANB3 behaves (**Figure 3B**). We then validated the previous report that RNR-α overexpression suppresses H-*ras*^G12V^-induced focus formation in NIH-3T3 fibroblasts (**Figure S9A**). Nuclear RNR-α, regardless of catalytic activity, also suppressed focus formation similar to that shown by RNR-α global overexpression (**Figure 3C**). Consistent with this observation, shIRBIT lines—that have elevated nuclear RNR-α levels^25^ (**Figure S1B**) but have elevated dNTP-producing capacity^39^—also showed limited ability to form foci (**Figure 3D**, top row). This outcome documents that nuclear RNR-α is likely the principal cause of RNR-α-overexpression-promoted tumor suppression and further indicates that inhibition of basal ZRANB3 activity could be the(a) root cause of the observed tumor suppression.

We also examined the effect of PCNA-overexpression. Overexpression of PCNA rescued the effect of RNR-α-promoted tumor suppression (**Figure 3C**, lower row), similar to what we observed for the nuclear-RNR-α—ZRANB3-interaction-driven DNA-synthesis inhibition. These data together predict that ZRANB3-knockdown would suppress focus formation in NIH-3T3 cells. We transfected our sh-ZRANB3 line with H-*ras*^G12V^ and measured the resulting number of foci relative to two different control lines. These data showed that ZRANB3-knockdown strongly reduced focus formation; RNR-α overexpression had little effect on the focus content in these cells (**Figure 3D**, lower row).

## Conclusion

Nucleos(t)ide drugs are a venerable class of antimetabolites that share numerous chemical similarities^2, 18^. Several of these molecules show no statistically-differentiable efficacy in “Phase IV” clinical trials^40^. However, there is certainly evidence that these drugs do behave differently: e.g., the clinical efficacy of CLA and F2C are quite divergent^41^; and of course, F2C shows efficacy against solid tumors that is not manifest by ClF, CLA, or FLUMP^2, 15^. Understanding the factors responsible for the variable mechanism(s) of action is of significant interest.

Unfortunately, gaining mechanistic insights is difficult because it is widely understood that nucleos(t)ide drugs are pleiotropic. From our work and that of others, it is clear that RNR-reductase activity inhibition is a key early step along with the multifactorial mode of action underpinning several nucleoside antimetabolites^2^. However, following RNR-α inhibition, other processes are critical^15, 16, 18, 19^. The importance of each independent or coordinated mechanism to drug efficacy is challenging to assess, and hence there could easily be many unknown factors contributing to drug efficacy and clinical applications that remain to be found. Characterizing additional modes of drug action is thus of both fundamental and translational relevance and could potentially lead to new avenues of therapeutic discovery/hint at the mode(s) of action of utilized, but unannotated, drugs.

We were able to score a common mode of action for CLA, ClF, and FLUMP that involves eliciting reductase-activity-downregulated RNR-α-hexamers. Critically, all of these compounds are analogs of the native nucleoside dA, the triphosphate of which (dATP) binds an allosteric site on RNR-α. This mode of action (both site selectivity and RNR-α-specific-hexamerization) is not shared by 3-AP and F2C^12^, which were used as comparisons in this study. Our data show that at least in the early time points, almost all of the DNA-synthesis inhibition for ClF, CLA, and FLUMP is directly associated with inhibition of ZRANB3. Indeed, in cells devoid of ZRANB3, some of these drugs increase DNA synthesis, as opposed to blocking it. Thus, the non-DNA-damage-associated activity of ZRANB3 is an early casualty of RNR-α-hexamer-inducer drugs.

Conversely, in the late stage of toxicity, shZRANB3 lines are similarly affected by nucleoside antimetabolites as in control lines. We believe that this is most likely because late-stage cytotoxicity is dominated by DNA-damage induction, futile damage/repair cycles, base-pair mis-incorporation, breakdown of mitochondria integrity and changes in cytosine methylation, among others^16, 18, 19, 42^. Under these conditions, PCNA is ubiquitinated^28–30^ rendering ZRANB3 less able to interact with RNR-α. This may, at first glance, suggest that RNR-α—ZRANB3 interaction is not critical for ClF action. However, we do not know which conditions—early or extreme end-point—reflects the mode of action of ClF/CLA/FLUMP in patients (but we do know that the early state is essential for drug action). Furthermore, there is some evidence that RNR-α-hexamer-inducers modulate DDR: ClF exerts larger effects on DDR than 5-fluorouracil^43^. Indeed, both FLUMP and ClF are inhibitors of the DDR, which could be explained by impairment of ZRANB3. Further investigations into the relationships we uncover could readily be targeted in this direction. Regardless, our data imply that the nuclear RNR-α—ZRANB3 axis has been inadvertently tapped into by these FDA-approved drugs. Our study is thus a starting point to highlight that this action may be pursuable in terms of a novel intervention.

We postulated that more insights could be gleaned from investigating the link between the ZRANB3— RNR-α axis and focus formation. We here disclose the novel finding that ZRANB3 is required for invasion and H-*ras*^G12V^-induced focus formation. We thus implicate the RNR-α-mediated ZRANB3-inhibition as the cause of RNR-α-driven reduction of tumor foci, reported seminally by Wright et al.^23^, and later by other research groups^2^. Importantly, our data do not show that ZRANB3 is an oncogene; it is most likely that H-*ras*^G12V^-transformed cells are more reliant on ZRANB3 to promote/sustain growth. We thus consider this observation as an example of non-oncogene addiction^44^. Our data, in fact, are strongly consistent independent work showing that both heterozygous and knockout ZRANB3 mice are more prone to myc-induced tumorigenesis than wt mice^45^. Clearly, examples of non-oncogene addiction are ripe for anti-cancer drug discovery, rendering it likely that ZRANB3 is an unheralded cancer-drug target. We will thus in the future focus our efforts on developing small-molecule interventions that will mimic nuclear RNR-α and inhibit the DNA-synthesis-promoting activities of ZRANB3. We hope that others will similarly take up the baton in this area.

## Abbreviations

3-AP: triapine
CLA: cladribine
ClF: clofarabine
DPA: dual-pulse whole-nuclei staining assay
F2C: gemcitabine
FLUMP: fludarabine monophosphate
IF: immunofluorescence
PLA: proximity ligation assay
WB: western blot

## On-line Methods

### Focus formation and scratch/wound healing assays in NIH-3T3 cells

For focus formation assays, NIH-3T3 cells were transfected with H-*ras*^G12V^ (0.5 µg) and indicated transgenes and/or empty vector (2.5 µg) in a 6 well plate using lipofectamine LTX and plus reagent. After 2 days, cells were split and equal numbers of cells (0.5–0.7 million) were plated in a 6 cm dish. Cells were grown to confluence with medium exchanges every 2–3 days (after 4 days 4% serum media were used) for 2-3 weeks, until foci became visible, then cells were fixed in 4% cracked PFA for 30 min, then stained with crystal violet solution and destained with water as required. For scratch/wound healing assay, cells on glass plates were grown to confluence then scratched using a glass pipette to introduce a central linear scratch across the surface (two scratches were made perpendicular to each other and analyzed separately). 2D cell migration into the wound was measured by imaging on a Biotek Cytation III plate reader. The plot was generated by measuring the initial wound width and dividing the width of the wound as a function of time by the initial width for each set. Wound widths varied by less than 25% across all data sets.

### Imaging analysis

Cells were grown in 35 mm glass-bottom plates to appropriate confluence and treated as described. At the conclusion of the experiment cells were fixed by addition of 2 ml MeOH (–20 °C) for 20 min (– 20 °C). MeOH was removed and PBS was added and stored at 4 °C for days to weeks. NOTE: fixing with MeOH is critical for appropriate visualization of RNR-α. Cells were then washed 3 times with PBS and carried straight on to blocking (2 ml imaging blocking buffer: 3% BSA & 0.2% Triton x-100 in PBS at 37 °C for 1 h). Plates were washed 2 times with PBS (1 ml), then treated with primary antibody, in antibody incubation buffer: 1% BSA and 0.02% Triton x-100 in PBS, using 120–150 μl to cover only the glass portion of the plate. Plates were incubated at rt for 2 h. Primary antibody was removed and plates were washed two times with PBS (1 ml). Then secondary antibody (120–150 μl; 1:1000 dilution in antibody incubation buffer) was added (1 h). Plates were washed one time in PBS containing 1 μg/ml DAPI, then once with PBS and stored at 4 °C until ready to image.

Only for imaging BrdU/EdU-treated cells: directly before blocking, cells were treated with 1 M HCl at 4 °C for 10 min, then 2 M HCl at rt for 10 min, then warmed to 37 °C for 10 min. After this time cells were treated with PBS two times, then 50 mM HEPES pH 7.6. Cells then were treated with a freshly made mix of 1 mM CuSO_4_ (from a 100 mM CuSO_4_ stock in water), 10 mM sodium ascorbate (made as a 100 mM stock in 500 mM HEPES pH 7.6) all in 50 mM HEPES pH 7.6 and 15 μM Cy5-azide. Cells treated with this mixture were left for 20 min in the dark. Note: this mix must be made just before use. Then cells were treated with imaging blocking buffer for 1 h, then washed twice with PBS, then primary antibody buffer was applied and subsequent steps were carried out as described above, starting from the primary antibody addition step. Note Clone MoBu-1 anti-BrdU is essential for the DPA (dual pulse imaging assay).

PLA was carried out as suggested by Sigma Aldrich using the procedure for Doulink® in situ red starter kit with mouse/rabbit, primary antibody incubations were as described above.

### Antibodies

RNR-α [AB81085 (Rb), AbCam. 1:1000 (WB), 1:200 (IF); 60073-1-Ig (Ms), Proteintech, 1:2000 (WB), 1:200 (IF), 1:300 (PLA for both HeLa and U2OS cells)]; RNR-b [AB57653 (Ms), AbCam, 1:1000 (WB), 1:200 (IF)]; BrdU [NA61, Calbiochem, 1:1000 (Clone MoBu-1, DPA, selective for BrdU over EdU)]; ZRANB3 [A303-033-A (Rb), Bethyl Laboratories. 1:2500 (WB); 1:200 (IF); HPA035234 (Rb), ATLAS, 1:200 (WB for U2OS cells and IF for HEK293T cells), 1:300 (PLA for HeLa cells), 1:25 (IF and PLA for U2OS cells)]; goat anti-rabbit Alexa-488 [A11008, Invitrogen. 1:1000 (IF)]; donkey anti-rabbit Alexa 647 [Ab150075, AbCam, 1:1000 (IF)]; donkey anti-mouse Alexa 568 [Ab175472, AbCam, 1:1000 (IF)].

WB = western blot; IF = immunofluorescence; PLA proximity ligation assay. Ms = mouse. Rb = rabbit.

### siRNAs

siRNA was delivered using dharmafect as per the manufacturer’s protocol. Unless otherwise indicated in figure legends, cells were transfected at 25% confluence, and typically left for 2 days prior to experimentation. siZRANB3(1) siRNA was purchased from Dharmacon (D-010025-03-0005). siZRANB3(2) and siZRANB3(3) were purchased from Invitrogen (s38487 and s38488). Two negative control siRNAs were purchased from Santa Cruz Biotechnology (sc-37007 and sc-44230). The sequences of the 3 different ZRANB3-targeting siRNAs were as follows:

siZRANB3 or siZRANB3(1): GAUCAGACAUCACACGAUU
siZRANB3(2): CAAGAGAUAUCAUCGAUUAtt
siZRANB3(3): GAUUCGAUCUAAUAACAGUtt

Invitrogen designed their siRNAs by adding “tt” to the 3’-ends to improve the stability of the siRNA.

RNR-α siRNA (D-004270-01-0005, D004270-02-0002, and D004270-03-0002) and RNR-β siRNA (D-010379-05-0005) were purchased from Dharmacon.

The sequences of the RNR-α siRNAs were as follows:

siRNR-α or siRNR-α(1): GAACACAUACGACUUUA
siRNR-α(2): GGACUGGUCUUUGAUGUGU
siRNR-α(3): GCACAGAAAUAGUGGAGUA.

### shRNAs

The following shRNA sequences were used in pLKO1 vector. The shRNA plasmids for control were gifts from Dr. Andrew Grimson (Cornell University, Ithaca, NY). The shRNA plasmids for ZRANB3 (of which one was functional) and IRBIT were purchased from Sigma-Aldrich. For details, see lentivirus production and infection method sections.

shControls:

1. CGCGATCGTAATCACCCGAGT (lac Z);
2. GTCGAGCTGGACGGCGACGTA (GFP).

shIRBIT

1. CCAGAAAGTTGATTCTTTA (mouse)

shZRANB3

1. TGGTCTTTGCGCACCATTTA (mouse)

### Generation of knockdown cell lines using lentivirus

HEK293T packaging cells were seeded and grown overnight in antibiotic-free media in 6 well plates. At 80% confluence, each well was transfected with packaging plasmid (pCMV-R8.74psPAX2, 500 ng), envelope plasmid (pCMV-VSV-G, 50 ng) and pLKO vector (500 ng) using TransIT.LT1 as per the manufacturer’s protocol. After 18 h media were removed and replaced with 20% serum-containing media. After 24 h, media were collected, spun down and passed through a 0.45-micron filter and used directly.

Cells in log phase were treated with 1 ml of virus supernatant (from above) in 8 μg/ml polybrene in a total of 6 ml of media in a 6-well plate. After 24 h, media were removed and replaced with media containing 2 μg/ml puromycin (which was 100% toxic to all lines used in this study). Cells were cultured till plate was confluent, then cells were split and moved to a 10 cm dish in 2 μg/ml puromycin containing media and grown till confluent again. At this point, the line was considered to be “selected”, and expression of target gene was analyzed by western blot and compared to shRNA controls. Up to passage 5 was used for these cells and they were typically grown in 1.5 μg/ml puromycin.

### Chemicals and reagents

All chemicals were from Sigma-Aldrich and were of the highest grade unless otherwise stated. Cyanine5(Cy5)-azide was from Lumiprobe; ClF, CLA, and F2C were from AK Scientific. Fludarabine monophosphate (FLUMP) was from Selleckchem. TCEP, deoxyguanosine (dG), deoxyadenosine (dA), adenosine (A) and EdU (working concentration 20 μM) were from ChemImpex. Deoxycytidine was from Sigma-Aldrich. Triapine (3-AP) was synthesized in house as previously described. Hygromycin B was from Invitrogen. AlamarBlue^®^ was from Invitrogen, used as the manufacturer’s instructions. Minimal Essential Media, RPMI, Opti-MEM, Dulbecco’s PBS, 100X pyruvate (100 mM), 100X non-essential amino acids (11140-050) and 100X penicillin streptomycin (15140-122) were from Gibco. *Transfection*: Cells were transfected using Mirus 2020 (all small scale and growth/proliferation experiments, used according to manufacturer’s instructions) or Mirus TransIT.LT1 (for virus production). MycoGuard^®^ Mycoplasma PCR Detection Kit was from Genecopoeia. Cells were typically left for 24–48 h without changing media prior to harvest/assay. ECL substrate or ECL-Plus substrate were from Pierce and used as directed. Acrylamide, ammonium persulfate, TEMEDA, Precision Plus protein standard was from Bio-Rad. All lysates were quantified using the Bio-Rad Protein Assay (Bio-Rad), relative to BSA as a standard (Bio-Rad). PCR was carried out using Phusion Hot start II (Thermo Scientific) as per manufacturer’s requirements. All plasmid inserts were validated by sequencing at Cornell Biotechnology sequencing core facility. All sterile cell culture plastic-ware was from CellTreat, except for glass-bottomed dishes used for imaging that was from *In Vitro* Scientific. The cloning cylinders for harvesting cell colonies (3166-6) were from Corning. The PLA kit was purchased from Sigma Aldrich (Doulink® in situ red starter kit with mouse/rabbit).

### Equipment

Plate reader assays were carried out on a Biotek Cytation III plate reader.

All confocal imaging experiments except those involving U2OS cells were carried out on a Zeiss LSM710 confocal microscope (Cornell Biotechnology Imaging Core Facility), equipped with an Inverted Axio Observer.Z1 microscope with 405, 458, 488, 514, 543, 561, and 633 nm laser lines. All U2OS cell-imaging experiments were performed on a Zeiss LSM700 confocal microscope (EPFL Bioimaging and Optics Platform), equipped with an Inverted Axio Observer.Z1 microscope with 405, 488, 555, and 639 nm laser lines.

### AlamarBlue^®^ assays

*For NIH-3T3 cells*, 4000 cells were plated in clear 96-well plates (total volume 100 μl), then 100 μl 2X compound was added in complete growth media and cells were allowed to grow for 48 h. After this time, 100 μl media were removed and AlamarBlue^®^ (10 μl) was added and cells were maintained at 37 °C for 3-7 h. For HEK293T cells, 10 cm plates were transfected with the stated siRNA and Dharmafect1 for 12 h, then 4000 cells were split into every 96 well plates (total volume 100 μl), then 100 μl 2X compound was added in complete growth media and cells were allowed to grow for 2 days. At the appropriate time point, number of cells was measured using fluorescence as per the manufacturer’s protocol. Data shown are mean +/− standard deviation of at least 8 independent replicates.

### Generation of Flp-In T-REx™ HEK293

Flp-In T-REx HEK293 cells were cultured in MEM supplemented with 10% FBS, 1X pyruvate and non-essential amino acids, 100 µg/ml zeocin and 1X penicillin/streptomycin in a humidified atmosphere at 37 °C with 5% CO_2_. To generate the line, HEK293TREX cells were co-transfected with the POG44 vector encoding the Flp recombinase and pcDNA5/FRT/TO-RNR-α(D57N)-NLS-2XFlag using Mirus 2020 in 6 well plates. Two days after transfection, the cells were split and passaged into 10 cm dishes, then after 1.5 d the cells were changed to selective media containing 150 µg/mL Hygromycin B. After 3 weeks, single colonies had formed, and these were harvested using cloning cylinders. Individual clones were grown in 12 well plates and verified for expression using western blot or imaging. To induce expression, cells were exposed to 100 ng/mL tetracycline.

### Quantification and analysis

#### Sample size n for figures with quantification

n for imaging experiments represents the number of single cells or in a few instances, panels quantified from at least five fields of view from at least two plates.

n for AlamarBlue® assays represents the number of individual wells under identical experimental conditions.

## Acknowledgements

The authors declare no conflict of interest. Shared instrumentation, personnel, and supplies in this work are supported by: the Novartis Foundation for Medical-Biological Research, Swiss National Science Foundation (SNSF) Project Funding, and NIH New Innovator (1DP2GM114850) (to Y.A.); and the National Centers of Competence in Research (NCCR) Chemical Biology (to Y.Z.). We thank Dr. Yuan Fu for collaboration in plasmid construction. We thank Professors Massimo Lopes (University of Zurich) and David Cortez (Vanderbilt School of Medicine) for the U2OS cell lines used in this study.

## Supplementary Figure Legends

**Figure S1.**
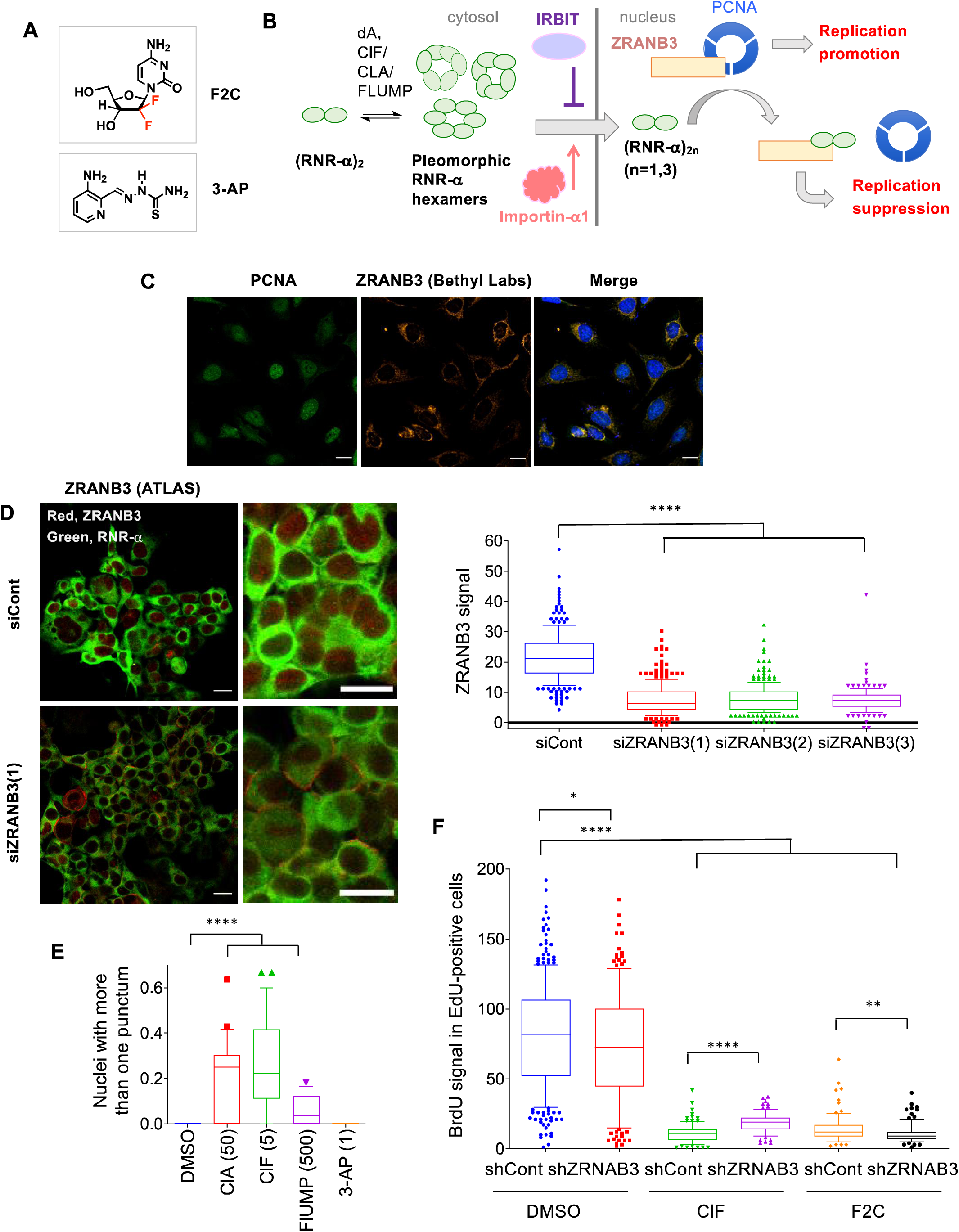
**A.** Structures of gemcitabine (F2C, RNR-α-subunit-targeting mechanism-based inactivator and non-RNR-α-specific hexamer inducer) and triapine (3-AP, RNR-β-subunit targeting inhibitor and non-RNR-α-hexamer inducer). **B.** The current model of dA(analog)-induced RNR-α nuclear translocation and downregulation of ZRANB3-dependent DNA-synthesis in non-DNA-damaged cells, featuring relevant interacting players. Briefly, phosphorylated derivatives of dA and indicated nucleoside drugs induce RNR-α-specific hexamerization. A proportion of these hexamers are nuclear-imported. Cytosolic proteins IRBIT and importin-α1 are negative and positive regulators respectively of this import process. Once inside the nucleus, RNR-α (irrespective of oligomeric state) disrupts PCNA—ZRANB3 binding which in non-stressed cells promotes DNA-synthesis. The resultant RNR-α—ZRANB3 interaction suppresses DNA-synthesis. **C.** Attempts to detect ZRANB3 by IF, using previously-employed anti-ZRANB3 antibody (Bethyl labs). HeLa cells were grown on glass-backed plates, fixed with MeOH, blocked and permeabilized, and indicated endogenous genes (PCNA and ZRANB3) were analyzed by IF. **D.** Similar to **B**, except that the ATLAS anti-ZRANB3 antibody was used to detect ZRANB3 in HEK293T cells treated with either siCont or 3 different siRNAs targeting ZRANB3. Representative images for siCont and siZRANB3(1) samples (left panel; the two columns correspond to low and high magnification images) and quantitation (right). **E.** Data from **Figure 1B** analyzed to show cells with two puncta or more. **F.** NIH-3T3 cells expressing either shControl or shZRANB3 were exposed to the different conditions (DMSO, ClF 5 µM, F2C 5 µM) for 30 min then dual-pulse labeling was performed. BrdU signal in EdU-positive cells was recorded. **Note**: In all figures here and elsewhere, unless otherwise indicated, siZRANB3 is equivalent to siZRANB3(1); siRNR-α is equivalent to siRNR-α(1). See Methods for sequence information.

**Figure S2.**
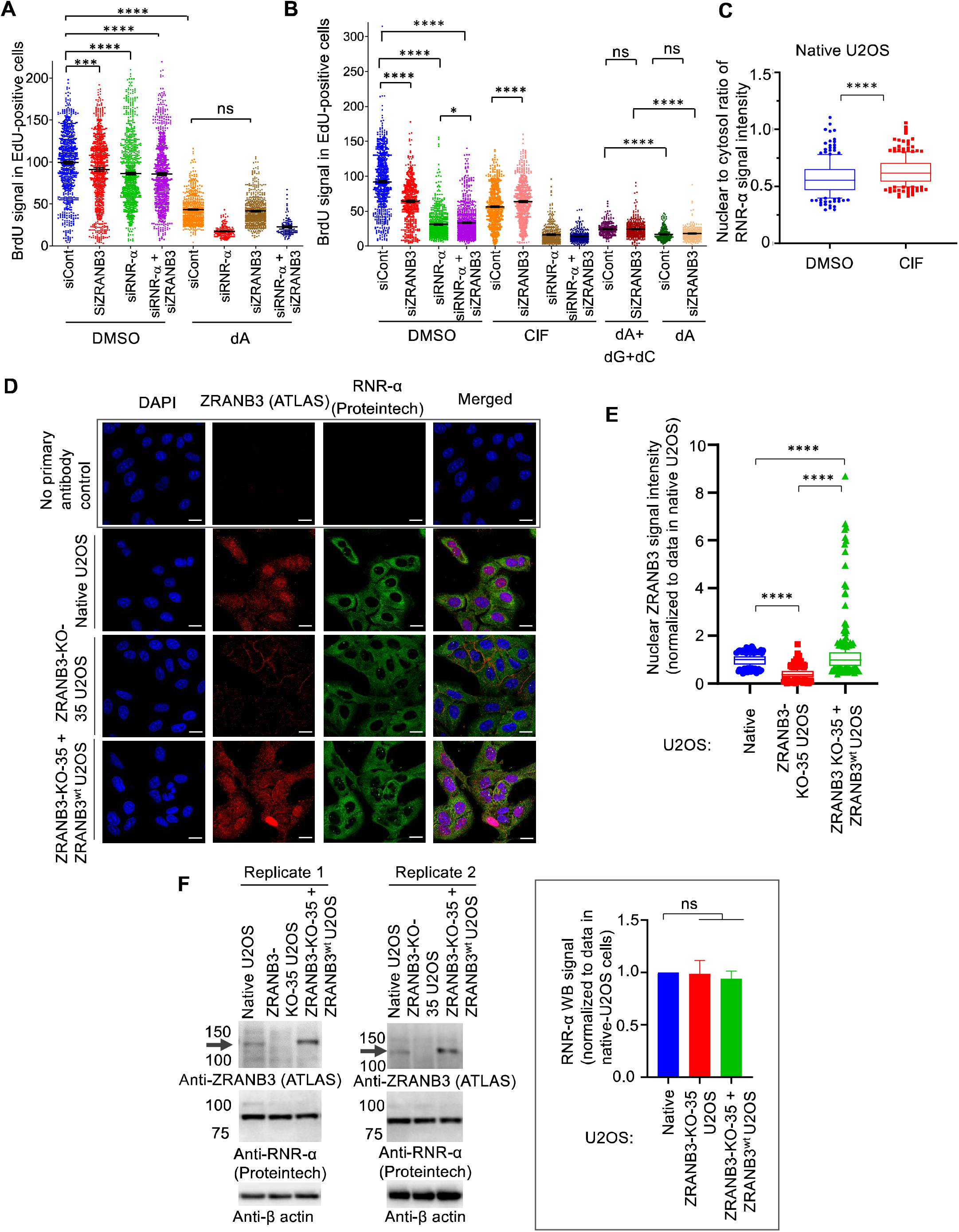
**A.** HEK293T cells were treated with the indicated 1:1 siRNA mixtures (where relevant, samples were balanced with siCont; whereas “siCont” samples contained 2X siCont) for 1 day followed by DMSO or dA (300 µM) for 30 min prior to undertaking dual-pulse labeling. BrdU signal in EdU-positive cells was analyzed using imaging. **B.** Similar to **A**, except siRNA treatment was administered for 2 days and nucleosides used were: ClF (5 µM), and dA (± dC, dG) (300 µM each). (Note: some of the data in this plot are recapitulated in **Figure 2A**). **C.** Quantitation of the IF data analyzing the extent of RNR-α-translocation in either ClF-(3 µM for 0.5-h) or DMSO-treated U2OS cells. **D.** U2OS, ZRANB3-KO-35 U2OS, and ZRANB3-KO-35 U2OS re-expressing ZRANB3^wt^ cells were fixed with −20 °C methanol prior to blocking, and incubation with primary antibodies, anti-ZRANB3 (ATLAS, 1:25) and anti-RNR-α (Proteintech, 1:200). After washing and applying the secondary antibodies, the cells were analyzed by confocal imaging. Top row: representative images showing the specificity of the secondary antibodies. Other rows: representative IF images of U2OS, ZRANB3-KO-35 U2OS, and ZRANB3-KO-35 U2OS re-expressing ZRANB3^wt^ in order from top to bottom. Scale bar, 10 µm. **E.** Quantification of data in **D**, reporting relative changes in nuclear-ZRANB3 protein levels in U2OS, ZRANB3-KO-35 U2OS, and ZRANB3-KO-35 U2OS re-expressing ZRANB3^wt^. **F.** Although validation of antibody-specificity via IF is of direct relevance to PLA (**Figure S3**) that requires IF-procedure, western-blot-based antibody-validation is also performed: indicated cell lines were lysed, total lysate proteins were normalized, and analyzed by western blots using the indicated antibodies. (Note: both anti-ZRANB3 and anti-RNR-α were used in performing PLA in **Figure S3**). **Note:** In ZRANB3-KO-35 U2OS re-expressing ZRANB3^wt^ (last lane), the re-expressed ZRANB3 bears both Flag- and HA-tags, hence the re-expressed protein appears to have a slightly higher molecular weight than the wild-type protein (arrow indicates the band corresponding to ZRANB3 molecular weight). Data from two independent biological replicates are shown. **Inset:** quantification of the western blot (WB) signal intensity corresponding to RNR-α.

**Figure S3.**
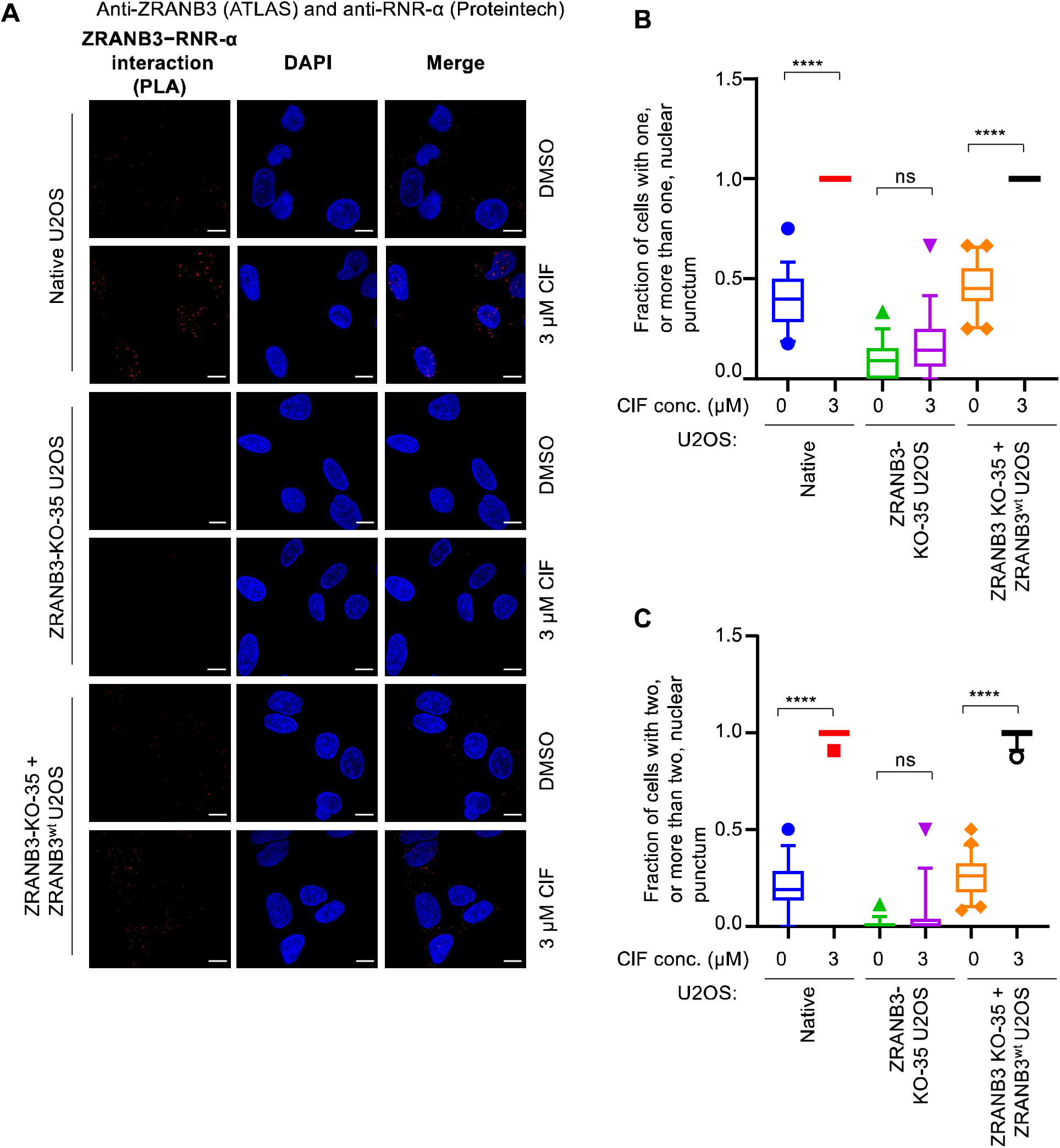
PLA in U2OS, ZRANB3-KO-35 U2OS, and ZRANB3-KO-35 U2OS re-expressing ZRANB3^wt^ validates that ClF promotes ZRANB3—RNR-α interaction in the nucleus. **A.** U2OS, ZRANB3-KO-35 U2OS, and ZRANB3-KO-35 U2OS re-expressing ZRANB3^wt^ were treated with either DMSO or 3 µM ClF for 30 min. Afterwards, cells were fixed with −20 °C methanol and PLA was performed following manufacturer (Sigma Aldrich)’s protocol using the primary antibodies: anti-ZRANB3 (ATLAS, 1:25) and anti-RNR-α (Proteintech, 1:300). Note: representative images are shown for each cell line either treated with DMSO (top) or ClF (bottom). Scale bar, 10 µm. **B.** Quantification of data from **A** reporting cells with one or more nuclear punctum. **C.** Quantification of data from **A** reporting cells with two or more nuclear puncta.

**Figure S4.**
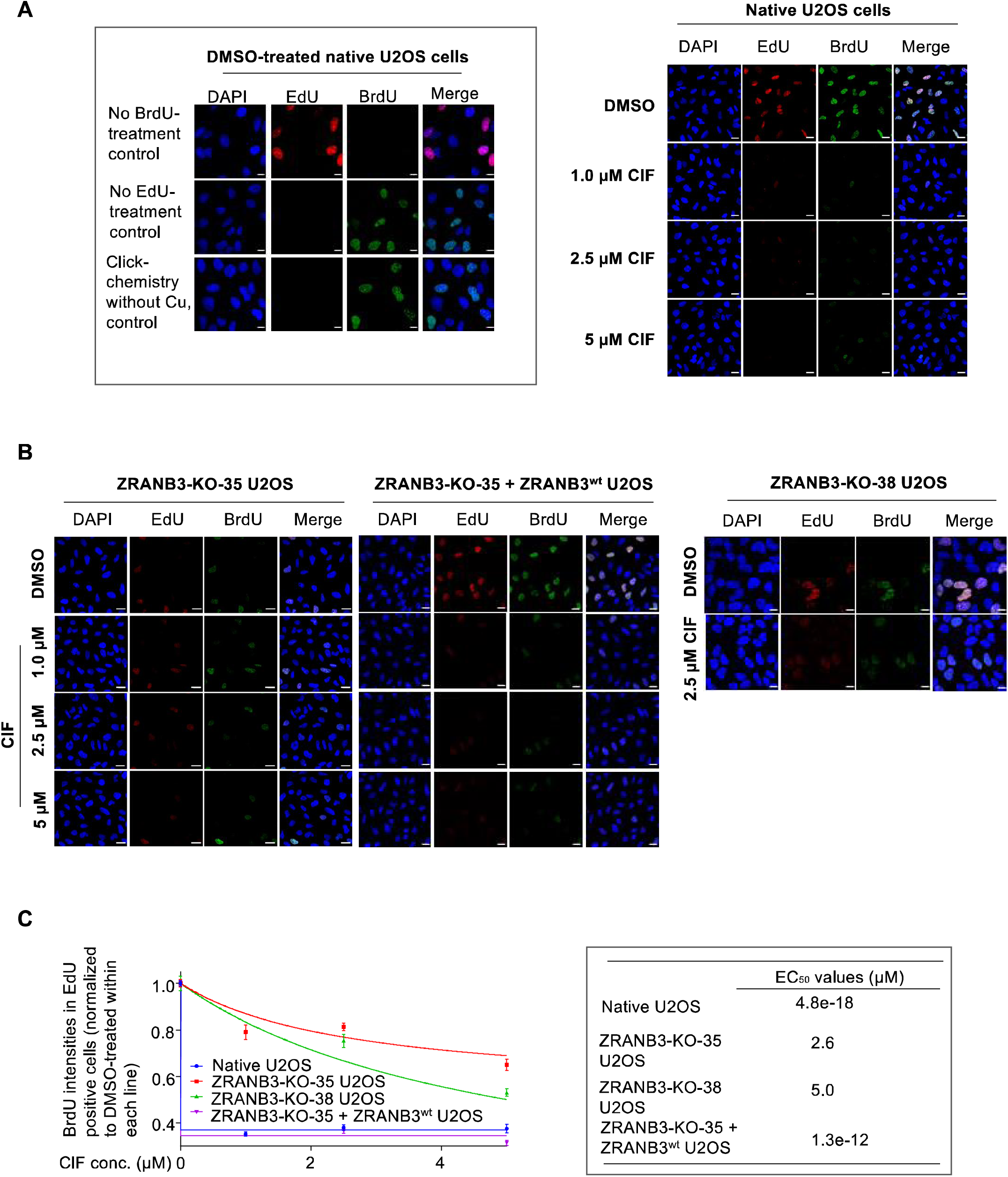
**A.** Representative images for **Figure 1F–G**. U2OS were treated with either DMSO or ClF (at indicated concentrations) for 30 min and subjected to DPA. Cells were washed with PBS, fixed with methanol, then analyzed by immunofluorescence imaging. Inset: representative images showing orthogonality of the EdU and BrdU detection methods [Click and anti-BrdU (Clone: MoBu-1, which does not detect EdU) antibody detection, respectively], and chromatic orthogonality of the dyes used (red, Cy5; and green, Alexa-488). Scale bar, 10 µm. **B.** Representative images for **Figure 1G**. ZRANB3-KO-35 U2OS, ZRANB3-KO-38 U2OS, and ZRANB3-KO-35 U2OS re-expressing ZRANB3^wt^, were subjected to treatment and analysis as in **A**. Scale bar, 10 µm. **C.** the quantification from **B** and **C** fit to the following equation y = {(1-k)/[1+(x/EC50)]} + k, where y is the normalized BrdU intensities in EdU-positive cells, x is the ClF concentration, and k is a constant representing the Y-axis value when [ClF] = ∞. Inset: EC_50_ values (µM) derived from the fit. Note: BrdU intensities in each line are normalized relative to that in the DMSO sample for each specific line (hence, all start at 1.0).

**Figure S5.**
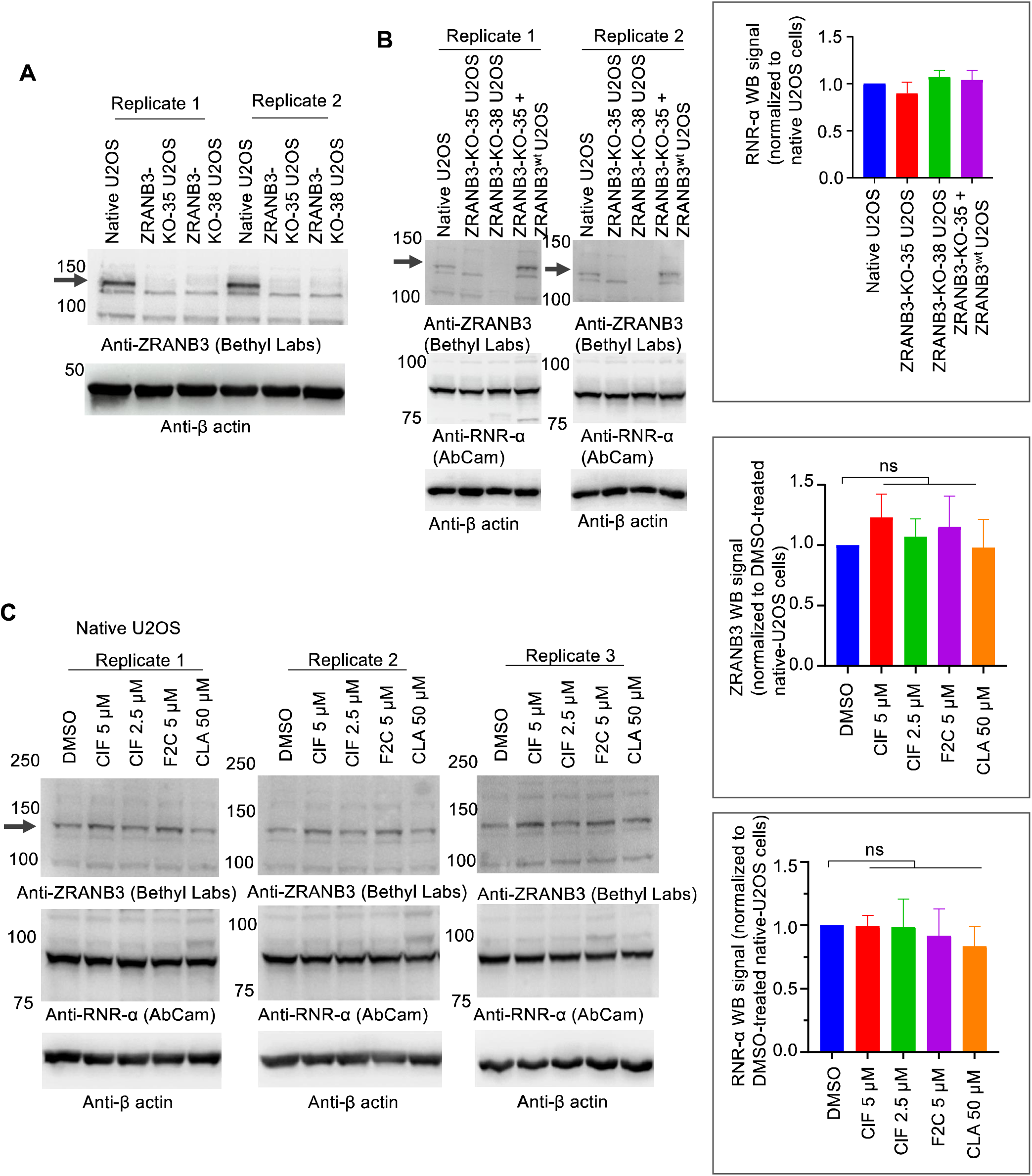
ZRANB3-knockout does not affect RNR-α expression and RNR-α and ZRANB3 expression levels are not affected by ClF, F2C, or ClA treatment in U2OS cells. An arrow denotes the band corresponding to the molecular weight of ZRANB3 in all sub-figures. **A.** Whole-cell lysates were normalized by Bradford assay, loaded onto a denaturing PAGE gel and analyzed by western blot using the indicated antibodies. Data from two independent biological replicates are shown. **B.** RNR-α protein expression level was not perturbed in ZRANB3-knockout U2OS cells or the same cells expressing full-length ZRANB3^wt^, compared to parent cells. Indicated cell lines were lysed, total lysate proteins were normalized, and analyzed by western blot using the indicated antibodies. (Note: in ZRANB3-KO-35 U2OS re-expressing ZRANB3^wt^ (the last lane), the re-expressed ZRANB3 bears both Flag- and HA-tags, hence the re-expressed protein appears to have a slightly higher molecular weight than the wild-type protein). Data from two independent biological replicates are shown. **Inset:** quantification of the western blot (WB) signal intensity corresponding to RNR-α. **C.** RNR-α and ZRANB3 expression levels are not affected by ClF, F2C or ClA (**Figure 1** and **S1A**) treatment in U2OS cells. The cells were treated with DMSO, ClF 5 µM, ClF 2.5 µM, F2C 5 µM, and CLA 50 µM for 30 min before harvest. The whole-cell lysates were normalized, and abundance of indicated proteins was analyzed by western blot analysis. Data from three independent biological replicates are shown. **Insets:** quantification.

**Figure S6.**
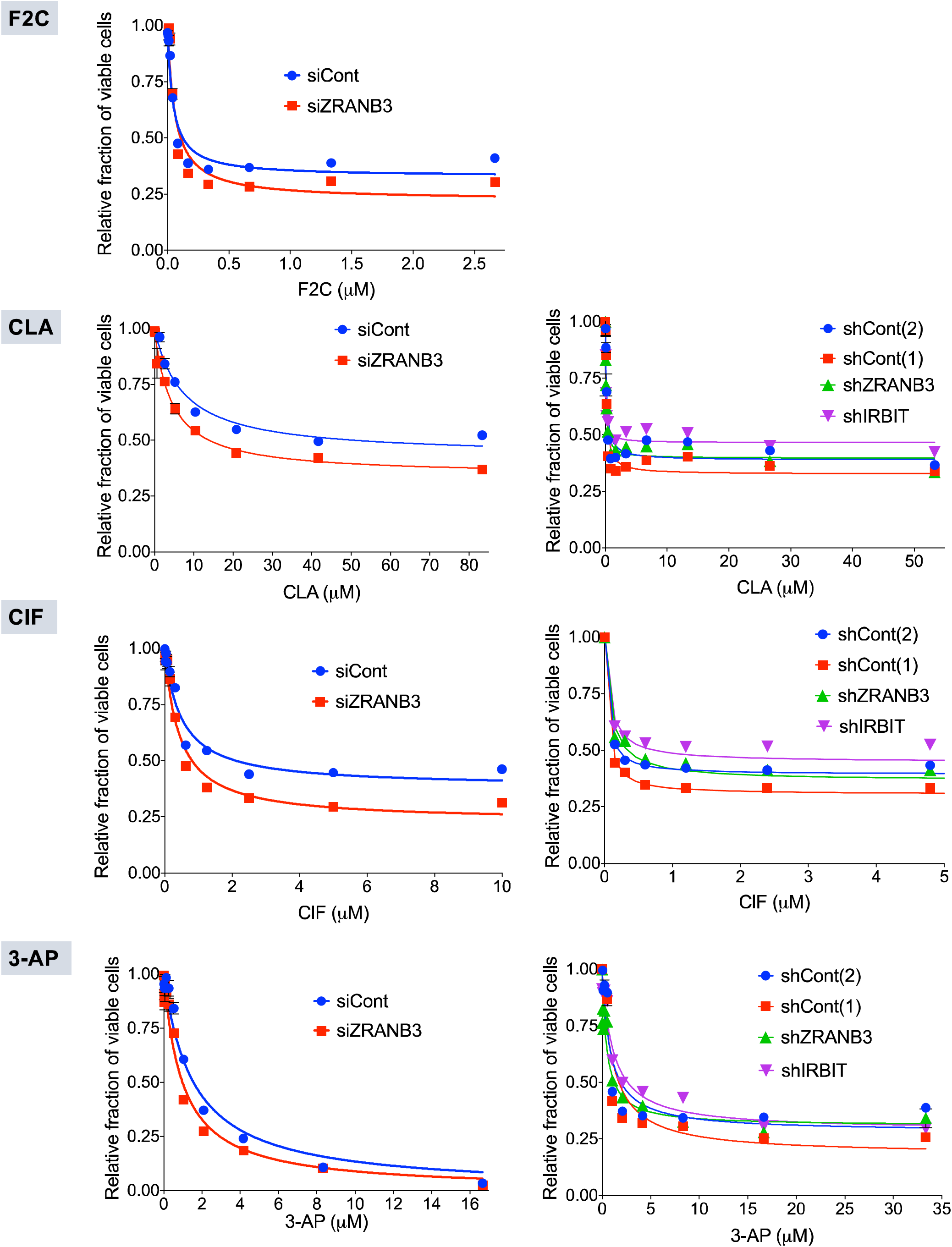
The indicated cell lines (<u>left panels</u>, HEK293T; <u>right panels</u>, NIH-3T3 wherein the indicated target gene was knocked down using either siRNA or shRNA, respectively) were treated with different doses of the indicated RNR-inhibiting drugs for 2 days. (See **Figure 1A** and **S1A** for the chemical structures of the drugs). For HEK293T cells, siRNA treatment occurred 12 h prior to treatment with the compound. After this time, the number of viable cells was measured using AlamarBlue®. Note: nucleoside antimetabolites at these concentrations elicit growth arrest and do not actually kill cells, thus curves do not go to zero. Data were fit to the following equation y = {(1-k)/[1+(x/EC50)]} + k, where y is the normalized BrdU intensities in EdU-positive cells, x is the ClF concentration, and k is a constant representing the Y-axis value when [ClF] = ∞. See also **Table 1**.

**Figure S7.**
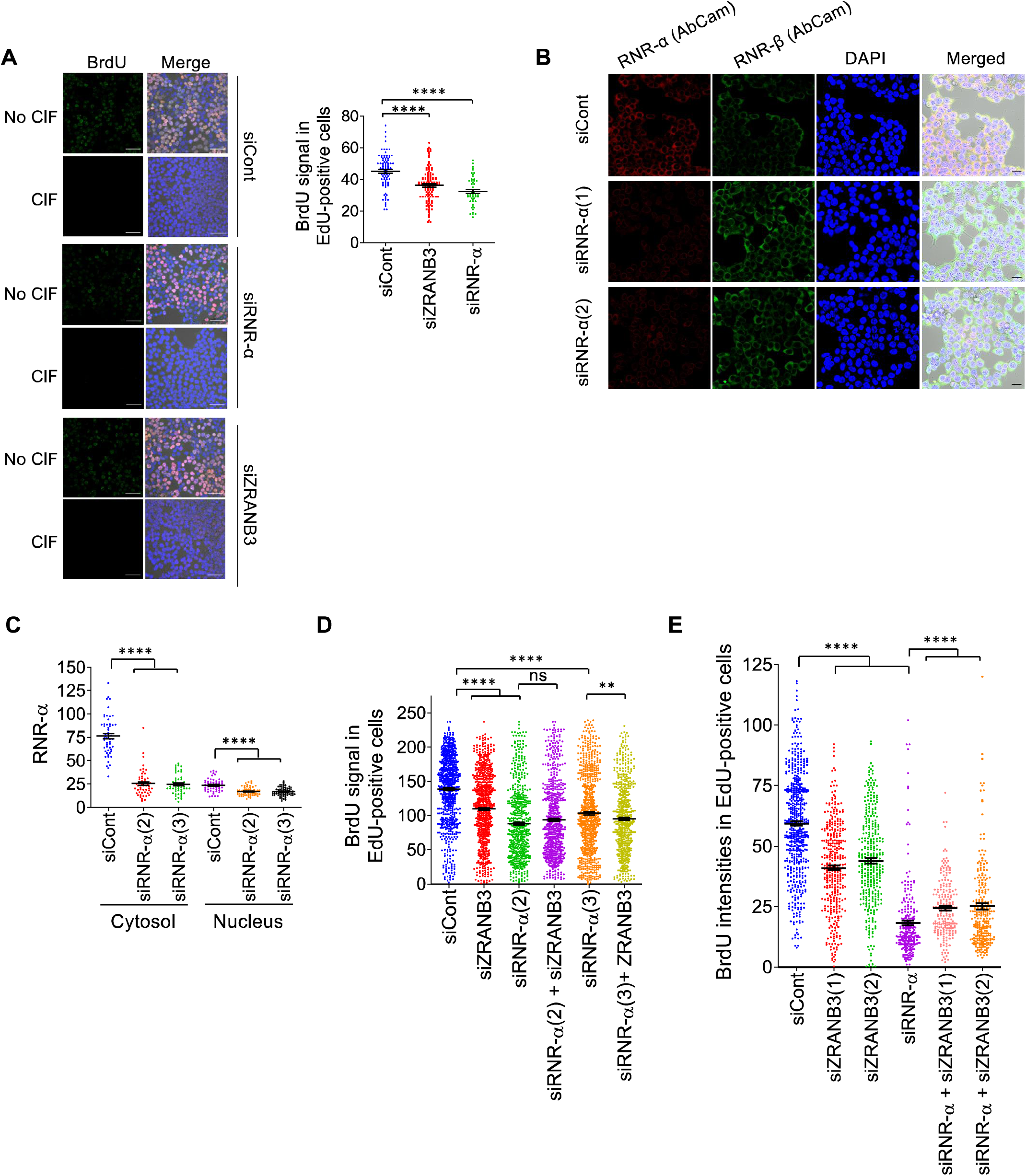
**A.** HEK293T cells were transfected with the indicated siRNA for 2 days after which time they were treated with DMSO or ClF (5 µM) for 2 h. After this time, dual-pulse imaging was carried out. Quantitation (plot on right) shows changes in basal BrdU incorporation in EdU-positive cells upon knockdown of the specific indicated genes in DMSO-treated cells. **B.** HEK293T cells were treated with the indicated siRNAs for 2 days and fixed with MeOH, stained with anti-RNR-α- and anti-RNR-β-antibodies, and imaged using a confocal microscope. **C.** Quantitation of images in **B**. **D–E**. HEK293T cells were transfected with the indicated siRNA combination for 2 days and DNA-synthesis rates were assessed by dual-pulse imaging. (*<u>Note 1</u>:* the same total siRNA amount was used for in each case, by balancing single siRNA doses with siCont). [*<u>Note 2</u>:* in **D**, although siZRANB3 decreases DNA-synthesis in siRNR-α(3)-treated cells, this is minimal compared to the effect on siCont cells].

**Figure S8.**
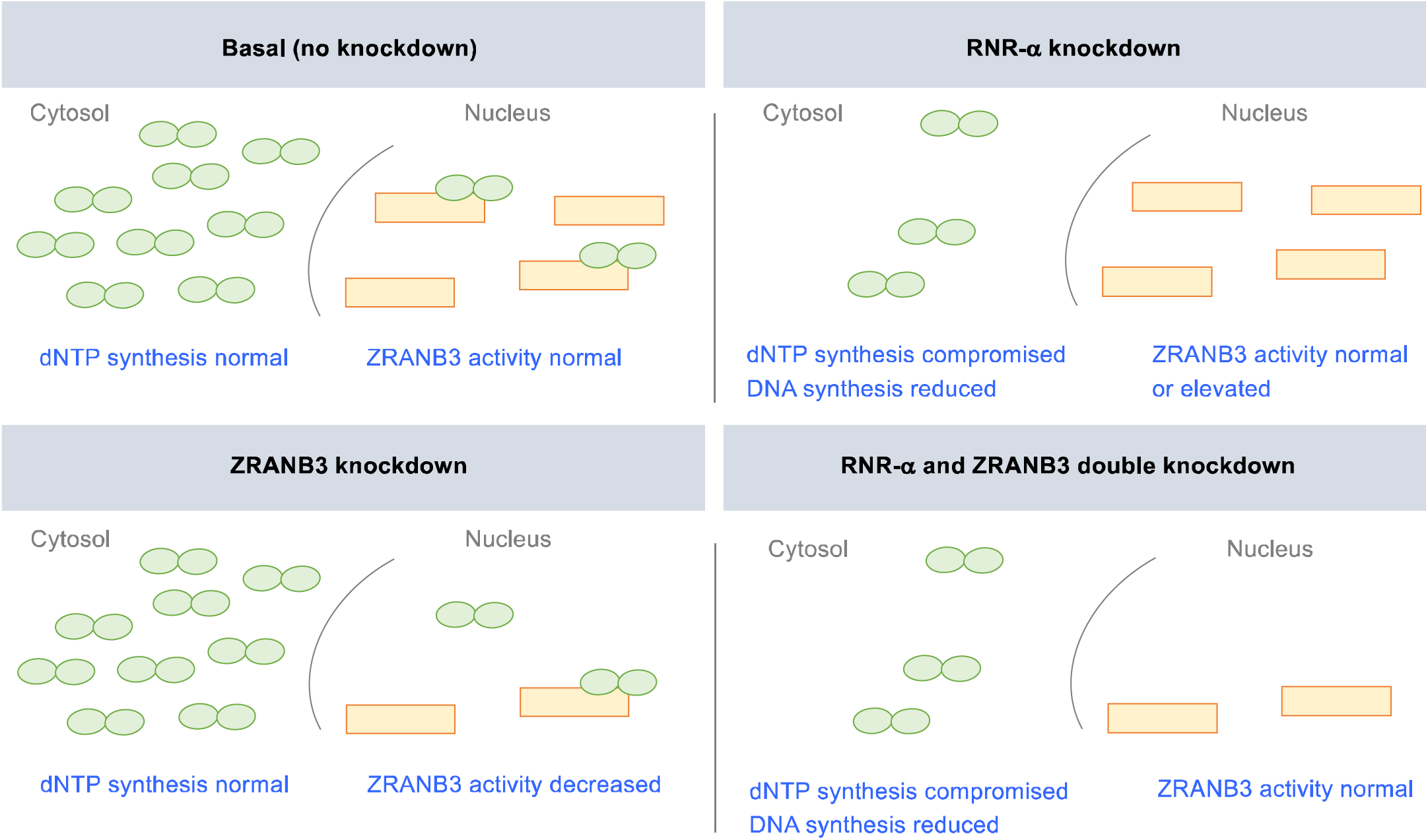
Model to understand the effects of different knockdowns on DNA synthesis. (The same color code for RNR-α (green ellipse) and ZRANB3 (yellow rectangle) is used as in **Figure S1B**. See also **Figure 1A**). *<u>Top</u> <u>left panel</u>*: normal. In the cytosol, RNR-α (together with RNR-β, not shown) provides dNTPs that are required for DNA synthesis. *<u>Top right panel</u>*: Knockdown of RNR-α thus decreases DNA synthesis by limiting dNTPs. RNR-α is also decreased in the nucleus, and hence active ZRANB3 is increased [or remain at similar level, if ZRANB3 is in excess under normal conditions (top left); however our data are consistent with ZRANB3 being limiting thus increase in active-ZRANB3 is expected]. *<u>Bottom left panel</u>*: In the nucleus, ZRANB3 is required for DNA synthesis. Knockdown of ZRANB3 decreases DNA synthesis (Recent report from another laboratory^45^, show that even loss of one ZRANB3 allele is phenotypic in some cases). *<u>Bottom right panel</u>*: When ZRANB3 and RNR-α are knocked down, there is decrease in DNA synthesis relative to basal (top left), but relative to RNR-α knockdown, there is no RNR-α to inhibit ZRANB3, so ZRANB3 knockdown has minimal effect.

**Figure S9.**
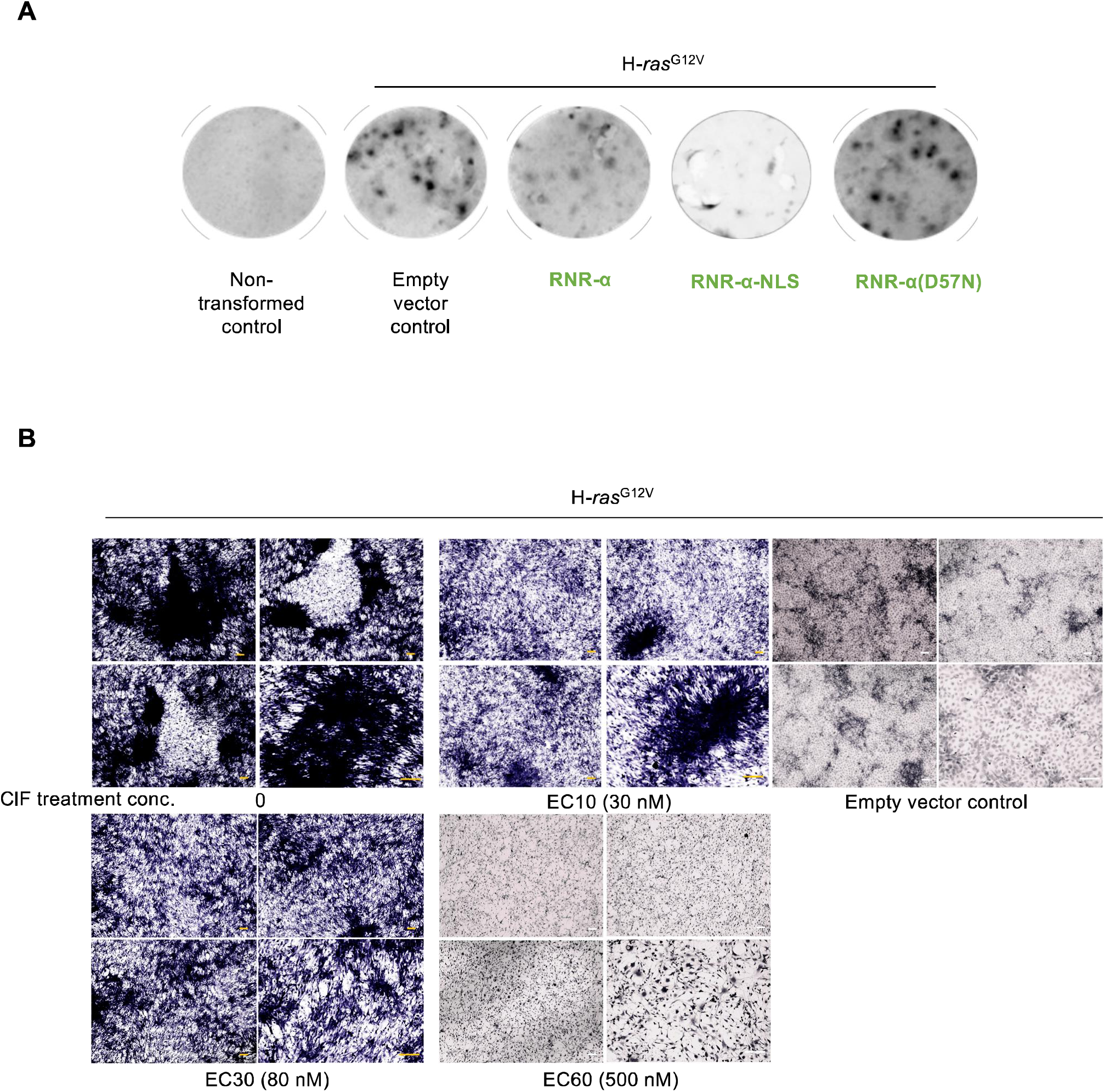
**A.** NIH-3T3 cells were transfected with the indicated plasmid combinations and then after 3 weeks, cells were stained with crystal violet, post washing, formaldehyde fixing, and staining. Also, see **Figure 3B–D**. **B.** Representative images from foci formation assays in the absence and presence of ClF treatment. See also **Figure 3B**. For each ClF concentration (i.e., the group of four images arranged in a square), three of the images (two on left hand side, and top right), were taken using a light microscope with a 4 X objective, and the image on the right on the bottom right was taken using a 10 X objective. Scale bar: 0.23 mm.

